# Actin retrograde flow actively aligns and orients ligand-engaged integrins in focal adhesions

**DOI:** 10.1101/071852

**Authors:** Vinay Swaminathan, Joseph Mathew Kalappurakkal, Shalin B. Mehta, Pontus Nordenfelt, Travis I. Moore, Koga Nobuyasu, David Baker, Rudolf Oldenbourg, Tomomi Tani, Satyajit Mayor, Timothy A. Springer, Clare M. Waterman

## Abstract

Integrins are transmembrane receptors that, upon activation, bind extracellular matrix (ECM) or cell surface ligands and link them to the actin cytoskeleton to mediate cell adhesion and migration^1,2^. One model for the structural transitions mediating integrin activation termed “the cytoskeletal force hypothesis” posits that force transmitted from the cytoskeleton to ligand-bound integrins acts as an allosteric stabilizer of the extended-open, high-affinity state^3^. Since cytoskeletal forces in migrating cells are generated by centripetal “retrograde flow” of F-actin from the cell leading edge, where integrin-based adhesions are initiated^4,5^, this model predicts that F-actin flow should align and orient activated, ligand-bound integrins in integrin-based adhesions. Here, polarization-sensitive fluorescence microscopy of GFP-αVβ3 integrin chimeras in migrating fibroblasts shows that integrins are aligned with respect to the axis of FAs and the direction of F-actin flow, and this alignment requires binding immobilized ligand and talin-mediated linkage to a flowing cytoskeleton. Polarization imaging and Rosetta modelling of chimeras engineered to orient GFP differentially with respect to the integrin headpiece suggest that ligand-bound αVβ3 integrin may be markedly tilted by the force of F-actin flow. These results show that actin cytoskeletal forces actively sculpt an anisotropic molecular scaffold in FAs that may underlie the ability of cells to sense directional ECM and physical cues.

---

To test the hypothesis that integrins are aligned in FAs by engagement to immobilized ligand and cytoskeletal flow, we employed mouse embryonic fibroblasts (MEF) that utilize αVβ3 integrins for adhesion and migration on fibronectin (FN). We co-expressed untagged β3 integrin together with GFP-αV integrin chimeras designed to impose different degrees of restraint on the mobility of GFP relative to integrin (Fig 1a, Fig S1). We then used polarization-sensitive fluorescence microscopy techniques that exploit the polarized excitation of and emission from the GFP transition dipole (henceforth referred to as dipole) to provide information about αVβ3 orientation and rotational dynamics^6,7^. The “αV-GFP-unconstrained” chimera had GFP fused via a 6 amino acid flexible linker to the C-terminus of the αV cytoplasmic tail, while “αV-GFP-constrained” was fused without the unstructured 5 amino acids at its N-terminus, in-frame to a β loop between Lys-259 and Asn-260 in the β-propeller of the αV extracellular headpiece (Fig 1a).

**Figure 1.**
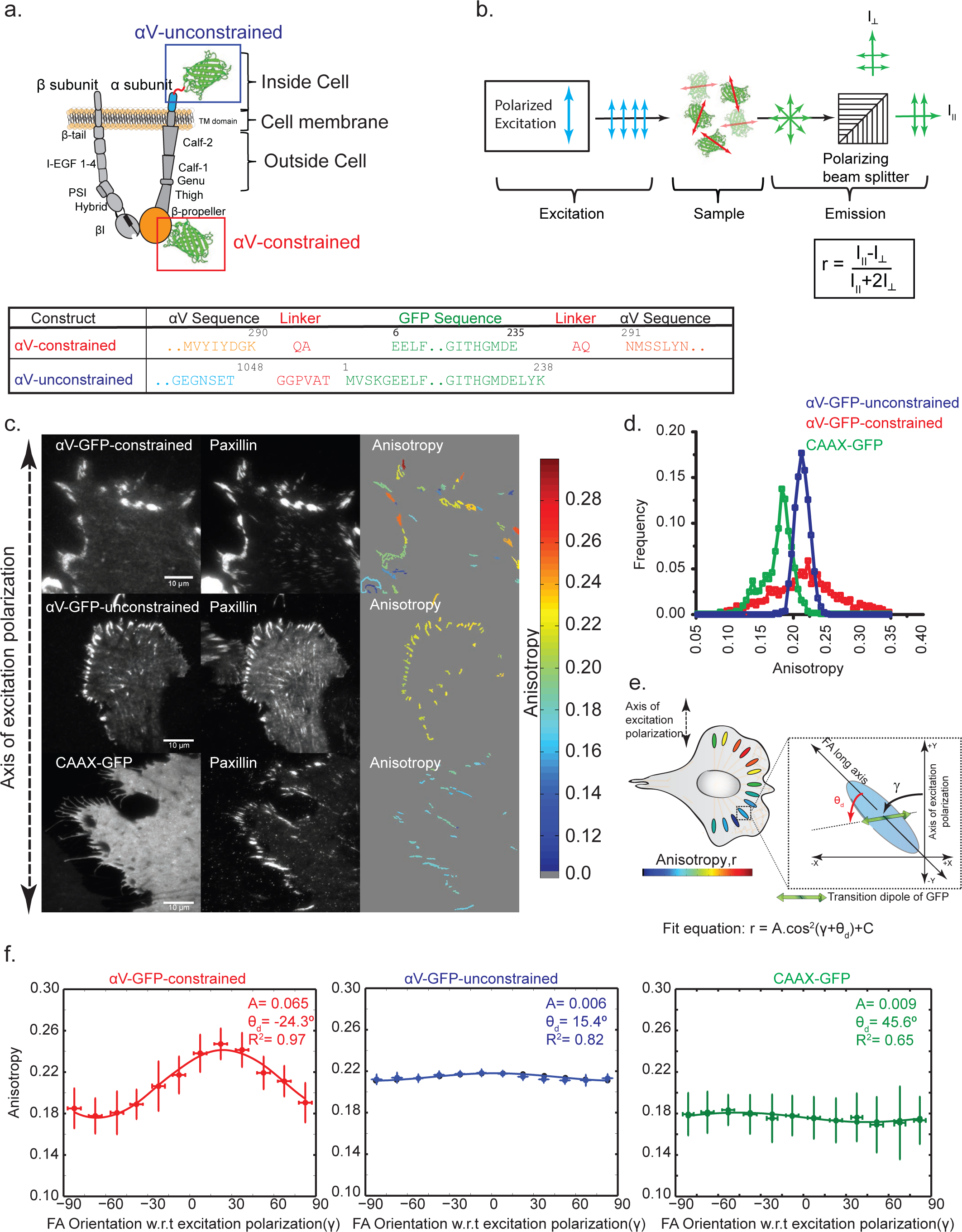
αV integrins are aligned and oriented relative to the FA long axis. **(a.)** (Top) Schematic diagram of αV-GFP constrained and unconstrained chimeras shown in the extended-open conformation of αVβ3 integrin. (Bottom) amino acid sequence and residue numbers at GFP-αV integrin junctions. **(b.)**(Top) Schematic of Emission Anisotropy-Total Internal Reflection Fluorescence Microscope (EA-TIRFM) setup and (bottom) equation defining anisotropy magnitude (*r*, I= intensity). **(c.)** (Left and center) Fluorescence micrographs of αVGFP-integrin chimeras (top left and middle left), GFP-CAAX (bottom left) or mApple-paxillin (center column) expressed in MEF plated on FN. Computed average emission anisotropy magnitude maps in segmented FAs (in the paxillin channel) from EA-TIRFM images of αV-GFP or GFP-CAAX (right panels). Orientation of excitation polarization with respect to the images (left), and anisotropy, keyed by color, (right). **(d.)** Histogram of emission anisotropy magnitudes of GFP chimeras in segmented FAs. **(e.)**Schematic diagram of the average emission anisotropy magnitude (keyed by color, bottom) for FAs of different orientations with respect to the excitation polarization axis in a cell for the case where the GFP transition dipoles are aligned with respect to the FA long axis as shown. Fit equation (bottom) where *r* is anisotropy; *C* is background; *A is* amplitude; γ is the angle of the FA with respect to the excitation polarization axis and θ_d_ is the orientation of the projection of the GFP dipole in the image plane with respect to the FA long axis**. (f.)** Plot of average emission anisotropy (r) in FAs vs FA orientation with respect to the excitation polarization axis (γ) for expressed GFP chimeras, overlaid with fit to the function in **(e).**Fit parameters shown in each plot. Error bar correspond to S.E.M, see Table I for n values for each experiment.

Cells were imaged by emission anisotropy total internal reflection fluorescence microscopy (EATIRFM), in which plane-polarized light excites those fluorophores in the specimen whose dipoles are oriented parallel to the plane of excitation (Fig 1b). The fluorescence anisotropy (*r*) of GFP emission is determined from the simultaneous collection of parallel and perpendicular components of the emission on separate cameras^10,11,12^ (Fig 1b), and is a measure of the co-alignment and rotational mobility of the GFPs that were excited by the polarized light^13^. EATIRFM imaging showed that for αV-GFP-unconstrained, emission anisotropy was generally low (Fig 1c,d), but was significantly higher in segmented FAs than regions outside FAs (Fig S2a). In contrast, either a GFP-tagged membrane-targeting sequence (CAAX-GFP) or soluble GFP exhibited a low value of emission anisotropy, irrespective of their localization to FAs (Fig 1c,d, Supp Fig S2a). Thus, even though GFP in αV-GFP-unconstrained is not constrained by linkage of both its N- and C-termini to integrin, its rotational mobility is limited within FAs.

We then performed EA-TIRFM analysis of αV-GFP-constrained. This showed that the average emission anisotropy in FAs was independent of expression level and FA size (Fig S2b), and was significantly higher than that for the unconstrained chimera or CAAX-GFP (Fig 1d. Fig S2a). However the average emission anisotropy values varied highly between individual FAs in a single cell, such that FAs with similar orientations with respect to the excitation polarization axis appeared to exhibit similar emission anisotropy values (Fig 1c right panel). This suggested that integrins may be co-aligned relative to the FA. To test this, we determined emission anisotropy in FAs from multiple cells, and binned FAs at 15° intervals of orientation relative to the excitation polarization axis (counter-clockwise being positive). This “radial sector analysis” confirmed that emission anisotropy varied as a function of FA orientation, with a peak emission anisotropy value at ~15-30°, and a minimum at ~75-90° (Fig 1f). We fit the emission anisotropy versus FA orientation data to a cos^2^ function that is expected if the dipoles have a fixed orientation in each FA with respect to the FA long axis (Fig 1e):

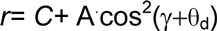

where *C* is the isotropic background; *A*, the amplitude, is related to the extent of GFP dipole co-alignment; γ is the angle of the FA with respect to the excitation polarization axis, θ_d_is the angle of the GFP dipole with respect to the FA long axis, and therefore γ+ θ_d_ is the angle of the GFP dipole with respect to the excitation polarization axis. The data for αV-GFP-constrained was well-fit by the cos^2^ function (R^2^=0.98) and the GFP dipoles on αV-GFP-constrained were aligned (A=0.070±0.01) at an angle of θ_d_ =-19.0 ± 9.8° (negative sign indicating clockwise with respect to the FA long axis (Fig 1f, Supp Table 1). In contrast, similar analysis for αV-GFP-unconstrained or CAAX-GFP showed poorer fits to the function and lower amplitudes (Fig 1f, Table 1). The orientation of the dipole with respect to the FA axis determined by this method was confirmed (-19.5°±3.85°) using Instantaneous FluoPolScope (Fig 4f top, Fig S5, a manuscript describing the method that is to be published elsewhere is attached), in which the dipole orientation in each pixel is measured directly (Fig S5). Thus, αVβ3 integrins are organized in an anisotropic fashion in FAs, with the dipoles in αV-GFP-constrained co-aligned with one another and specifically oriented relative to the FA long axis.

We next sought to determine the roles of ligand binding, ligand immobilization and activation in the alignment of integrins in FAs. To test the role of ligand binding, we plated and fixed cells co-expressing αV-GFP-constrained and mApple-paxillin on poly-L-lysine (PLL), while the requirement for ligand immobilization was tested by plating on supported lipid bilayers functionalized with RGD peptide that allows free ligand diffusion^14^. To determine if priming the integrin for ligand binding was sufficient for integrin alignment, cells were treated with Mn^2+^ and plated on PLL^15,16^. EA-TIRFM and radial sector analysis of cells under these conditions (Fig 2a, b) showed that compared to controls on FN, emission anisotropy was lower in integrin clusters, and the emission anisotropy *vs* FA orientation data function showed a poorer fit to the cos^2^ function and a lower extent of dipole alignment (A) under all conditions (Supp Table 1). Thus, binding to immobilized ligand promotes integrin co-alignment in FAs, but priming the molecule for high-affinity ligand binding is not sufficient for this alignment.

**Figure 2.**
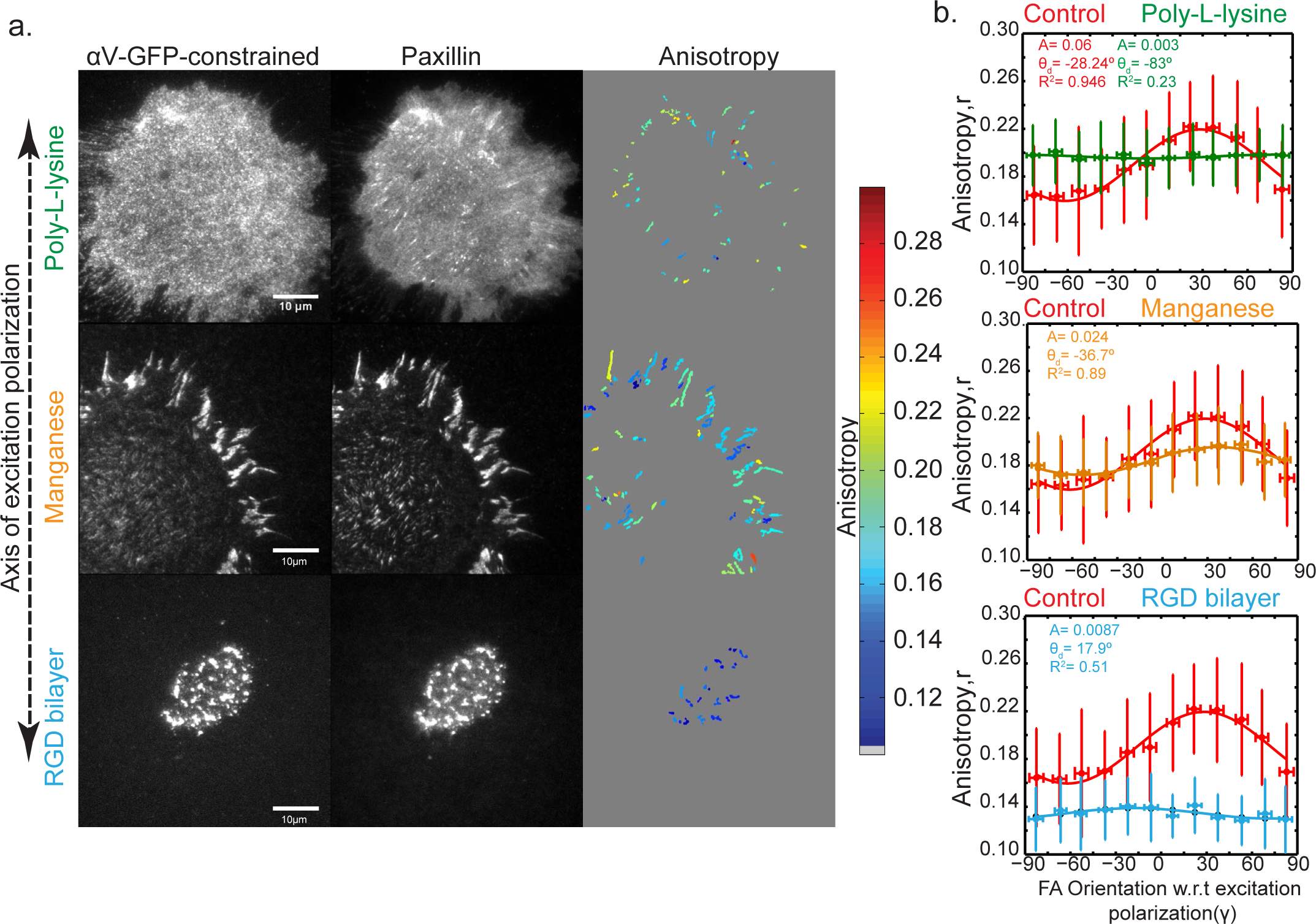
αV integrin alignment in FAs requires binding to immobilized ligand. **(a.)** Fluorescence micrographs of MEF co-expressing αV-GFP-constrained (left column) and mApple-paxillin (center column) and average emission anisotropy magnitude map computed from EA-TIRFM images of αV-GFP-constrained in segmented FA (right column), with orientation of excitation polarization relative to the images (left), and anisotropy, keyed by color, (right). Cells were plated on Poly-L-lysine (top), pretreated with 1mM Mn^+2^and plated on Poly-L-lysine (middle) or plated on RGD peptide conjugated to a supported lipid bilayer (bottom). **(b.)**Plot of average emission anisotropy (r) in FAs vs FA orientation (γ) with respect to the excitation polarization axis for αV-GFP-constrained, overlaid with fit to the function in Figure 1e, for the conditions shown in **(a.)**Fitting parameters shown in each plot. Error bar corresponds to S.D., see Table I for n values for each experiment.

We next sought to determine the role of F-actin organization and dynamics in integrin alignment in FA. We first asked whether the alignments of F-actin and integrins in FA were related. EATIRFM followed by radial sector analysis in segmented FAs of cells co-expressing αV-GFP-constrained and F-actin stained with Alexa-568 phalloidin showed that the emission anisotropy of F-actin within FAs varied as a function of FA orientation (Fig 3a, b). This data was well-fit to the cos^2^ function, with greater dipole alignment for F-actin than αV-GFP-constrained, although the orientation of the respective fluorophore dipoles differed with respect to the FA long axis (Fig 3b, Supp Table 1)^17^. Plotting the emission anisotropy of fluorescent phalloidin vs that of αVGFP-constrained in FAs showed a strong correlation (Fig S3a). Thus, αV integrin and F-actin both align at FA.

**Figure 3.**
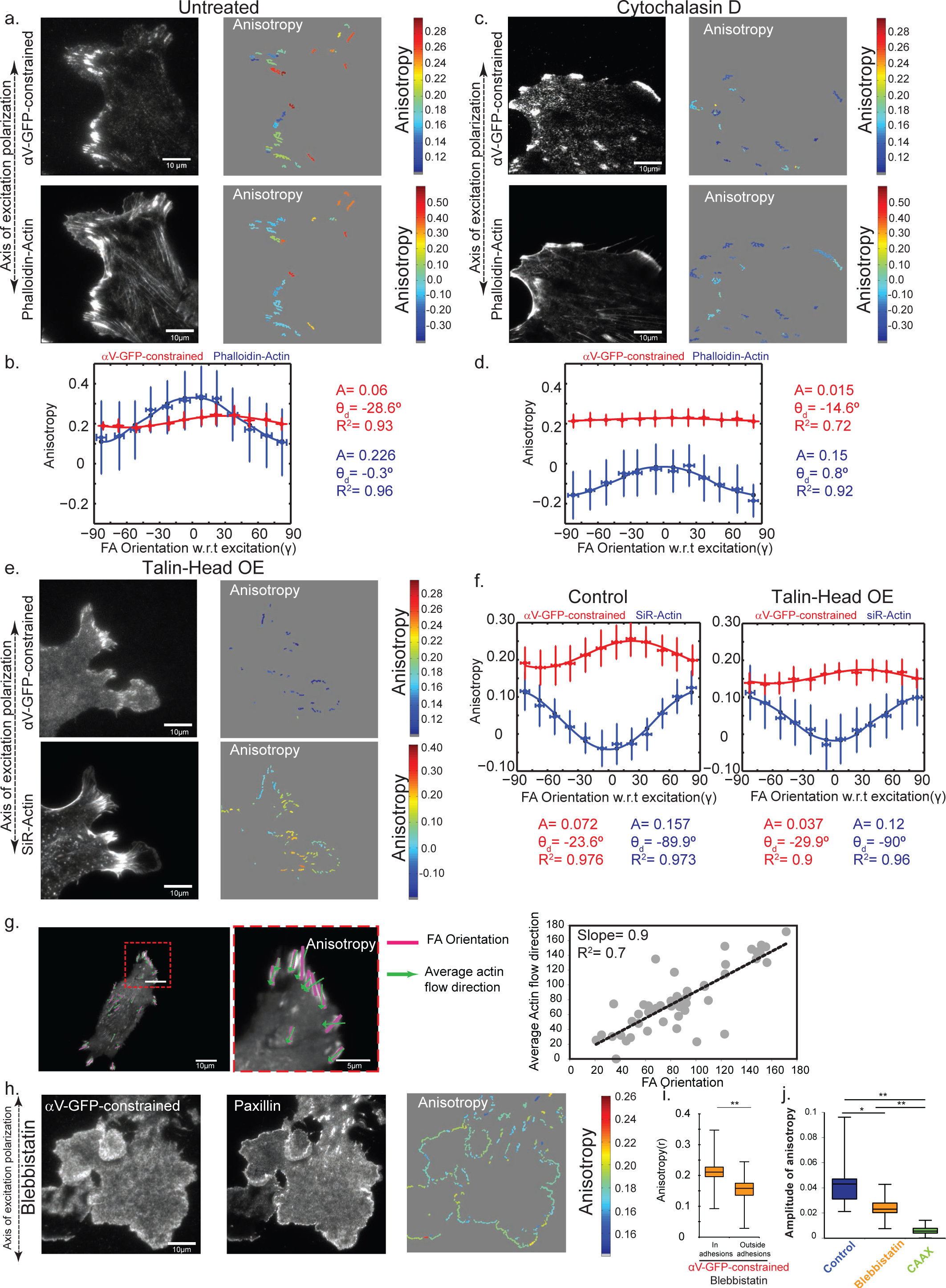
αV integrin alignment in FAs requires talin-mediated linkage to a flowing F-actin cytoskeleton. **(a,c,e,h)** Fluorescence micrographs of MEF expressing αV-GFP-constrained and F-actin stained with Alexa-568 Phalloidin **(a,c)**, or SiR 655-actin **(e)**, or co-expressing αV-GFP-constrained and mCherry-paxillin **(h)**. Corresponding anisotropy magnitude maps in segmented FA computed from EA-TIRFM images, with orientation of excitation polarization relative to the images denoted at left and anisotropy,*r*, keyed by color, (right). Scale bars= 10µm. **(b, d, f)**Plots of average emission anisotropy (r, n>300 FA) in FAs *vs* FA orientation (γ) with respect to the excitation polarization axis for expressed αV-GFP-constrained (red) and Alexa-568 Phalloidin ( **b, d,**blue) or SiR 655-actin (**f,** blue), overlaid with fit to the function in Figure 1e. Fit values for parameters shown in each plot. Error bars correspond to S.D., see Table I for n values for each experiment. **(c, d)**Cells were treated with 500nM cytochalasin-D. **(e, f)**Cells were over-expressing mcherry-tagged talin-head domain (Talin Head OE). **(g)** (Left) Fluorescence micrograph of expressed mApple-paxillin overlaid with the FA long axis (purple) and the average actin flow vector (green), obtained from quantitative fluorescent speckle microscopy analysis of expressed eGFP-actin (not shown). (center) Zoom of red box in middle panel. (right) Plot of FA orientation and average actin flow direction in segmented FAs (grey points represent n= 50 individual FAs from n= 5 cells). Scale in degrees relative to the y axis of the microscope, black line indicates linear fit of data. **(h, i, j)**Cells treated with 50µM blebbistatin. **(i.)** Box plot of average emission anisotropy of αV-GFP-constrained in nascent adhesions near the cell edge (In adhesions) or in the cell interior (Outside adhesions) **(j.)**Box plot of the amplitude (A) of the emission anisotropy from the cos^2^ fit of emission anisotropy *vs* orientation of a vector normal to the closest cell edge (see Fig S4A, Methods) on a pixel-by-pixel basis for αV-GFP-constrained in control (blue) or blebbistatin-treated cells (orange) or for GFPCAAX (green). In the box plots in (i) and (j), box includes 1^st^ quartile to 3^rd^ quartile, bar median, Whisker extends from minimum to maximum observation. *, p<0.001; **, p<0.0001, Kruskal-Wallis H test, see Table I for n values.

We then tested the requirements for an intact F-actin cytoskeleton and the talin-mediated linkage between F-actin and integrins in promoting integrin co-alignment in FAs. F-actin assembly was blocked with cytochalasin-D (Fig 3c,d), while integrin-actin link was inhibited by over-expressing mCherry-tagged talin-head domain^18,19^ (Fig 3e,f). SiR-655-actin^20^, which binds filaments but has its dipole oriented differently with respect to the actin filament axis than that of fluorescent phalloidin (Fig 3f, and S3), allows simultaneous imaging of αV-GFP-constrained, talin-head-mCherry and F-actin. Phalloidin or SiR-actin staining and EA-TIRFM radial sector analysis of F-actin showed that cytochalasin treatment disrupted the cytoskeleton and reduced F-actin emission anisotropy and its dependence on FA orientation (Fig 3d, Fig S3a, Table 1), while talin-head had little effect on F-actin morphology or its emission anisotropy (Fig 3f). In spite of these different effects on F-actin, radial sector analysis of αV-GFP-constrained showed that both treatments attenuated integrin emission anisotropy in FA and its angular dependence (Fig 3d,f Supp Table 1). Together this shows that although integrins and F-actin both align at FAs, F-actin alignment is not sufficient for integrin alignment. Instead integrin alignment requires an intact cytoskeleton, and is enhanced by the talin-mediated linkage between integrin and Factin.

We then pursued the role of F-actin retrograde flow in integrin anisotropy. We first determined the direction of F-actin flow in FAs by co-expressing mApple-paxillin and low levels of GFP-actin and performed TIR-fluorescent speckle microscopy^21^. Flow-tracking algorithms^5,22^ showed that F-actin underwent fast retrograde flow in the lamellipodium and slower retrograde flow in the lamella and FAs, as expected^5,23^ (Fig 3g). Plotting FA long axis versus local F-actin flow orientations showed a strong correlation and a slope near unity, indicating that F-actin flows along the FA long axis (Fig 3g). Together with results above, this shows that integrin alignment is correlated with the direction of F-actin flow in FAs.

To test if F-actin retrograde flow was sufficient for integrin orientation, we utilized the myosin-II inhibitor blebbistatin, which abolishes myosin-II-driven flow in the lamella and FAs, but leaves actin polymerization-driven flow in the lamellipodium intact^24^ (Supp Movie 1). This allowed determination of a regional correlation between F-actin flow and integrin alignment. As expected, blebbistatin eliminated cell polarity and blocked FA elongation, leaving a rim of nascent adhesions (NA) in lamellipodia^25^ (Fig 3h). EA-TIRFM of αV-GFP-constrained in blebbistatin-treated cells revealed higher levels of emission anisotropy in NA where retrograde flow remains intact, but significantly lower levels in the cell interior where NAs are absent and retrograde flow is blocked^24^ (Fig 3i). Since the long axis of NAs could not be determined, we analyzed emission anisotropy in NAs relative to the orientation of their closest leading edge (Fig S4). This emission anisotropy *vs* cell edge orientation data was well-fit to the cos^2^ function, albeit with significantly lower amplitude than control (Fig 3j, Fig S4b). Therefore, integrin alignment spatially correlates with F-actin retrograde flow, suggesting that flow is sufficient for integrin alignment in NAs. We thus conclude that integrin co-alignment in FA requires binding to immobilized ligand and is promoted by a talin-mediated linkage to a flowing actin cytoskeleton, and the direction of flow correlates with the direction of alignment.

We finally sought to model the orientation of integrins in FAs on the cell surface. This required estimating the orientation of the GFP dipole with respect to αVβ3 in our chimera, then placing the chimera into the frame-of-reference of the microscope with the dipole in the chimera oriented to match our experimentally determined GFP dipole orientation in FAs. Because the experimentally determined orientation is a 2D projection on the image plane of a dipole oriented in 3D, multiple orientations of αV-GFP-constrained could give the same 2D projection of the dipole orientation in FAs. To rule out some possible orientations, we thus determined the GFP dipole orientation in the FA of a second chimera in which GFP was positioned differently relative to integrin than that in αV-GFP-constrained. We generated “αV-GFP-less-constrained” by retaining the C- and N-terminal portions of GFP that show disorder in crystal structures (Fig 4a) ^8,11^. EA-TIRFM and radial sector analysis of cells expressing αV-GFP-less-constrained showed that the emission anisotropy *vs* FA orientation data was well-fit to the cos^2^ function, albeit with a reduced amplitude compared to that of αV-GFP-constrained (Fig 4 c,d, Supp Table 1), as expected from the extra residues at the integrin-GFP junctions. Importantly, the dipole was oriented at θ_d_= −87.7°± 3.6° with respect to the FA long axis, distinct from the θ_d_ =-19.0 ± 9.8° measured for αV-GFP-constrained. The orientation of the dipole in αV-GFP-less-constrained with respect to the FA long axis was confirmed (-85.5° ± 6.7°) using Instantaneous FluoPolScope (Fig 4f, Fig S5b).

**Figure 4.**
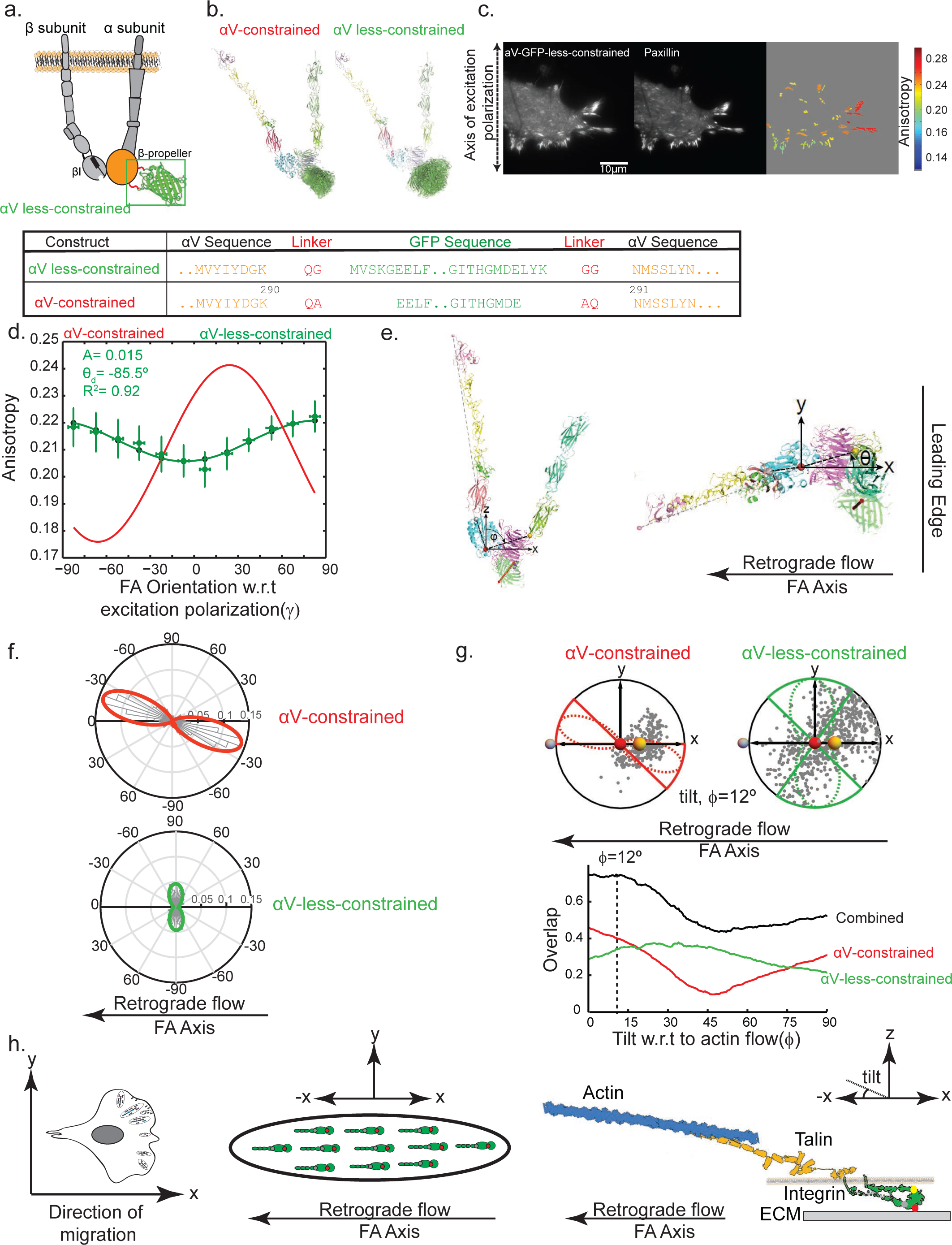
αVβ3 integrin is predicted to have the legs tilted in the direction of retrograde flow. **(a.)** (Top left) Schematic diagram of αV-GFP-less-constrained chimera in extended-open αVβ3 integrin. (Bottom) amino acid sequences and residue numbers at GFP-αV integrin junctions. **(b.)** Ribbon diagrams of ensembles of (left) αv-GFP-constrained or (right) αv-GFP-less-constrained chimeras. Representative, equal numbers of ensemble members output by Rosetta are shown superimposed on the integrin moiety. **(c.)** Fluorescence micrographs of MEF expressing αVGFP-less-constrained (left) and mCherry-paxillin (middle), and corresponding average emission anisotropy magnitude maps in segmented FA (right), computed from EA-TIRFM images, with orientation of excitation polarization relative to the images denoted at left and anisotropy, keyed by color (right). Scale bar= 10µm. **(d.)**Plot of average emission anisotropy (r) in FAs *vs* FA orientation (γ) with respect to the excitation polarization axis for αV-GFP-constrained (red) and αV-GFP-less-constrained (green), overlaid with fit to the function in Figure 1e. Fitting parameters shown in plot. Error bars correspond to S.D., see Table I for n values for each experiment. **(e.)** Ribbon diagrams of the extended-open conformation of αVβ3 integrin and GFP in 3V-GFP-constrained in the integrin/microscope frame of reference. Orientation of the GFP transition dipole (red double-headed arrow) is defined by a vector drawn from the N atom of Val^112^ to the O atom of Ser^147^. The integrin frame of reference plane is depicted by spheres at 3 Cα atoms (Asp of the RGD ligand, red), junction between the headpiece and the α3 leg (Arg^4380^, yellow) and junction between the headpiece and the β3 leg (Pro^111^, gray). Views along the microscope Y-axis (in the membrane plane, left) and z axis (membrane-normal, optical axis, right) shown. The integrin headpiece is oriented (-Z) toward the substrate, the ligand-point (red sphere) is fixed at the origin, the αV-leg (yellow sphere) and *β*-leg (gray sphere) junction points are within the X-Z plane, the cell leading edge is towards +X, the FA long axis is aligned along X, and F-actin flows from +X to –X. ϕ is the angle mage by the line between the αV-leg-junction and the origin with respect to the Z-axis and Θ is the angle between the projection of the origin-αV-leg-junction line in the X-Y plane and the Z-axis. to the origin with respect to the X-Z and XY planes of the microscope, respectively. In the reference state, ϕ=90° and Θ=0°. **(f.)** Polar plots of the orientations (scaled by polarization factor, defined on x-axis) of GFP dipoles for αVGFP-constrained (top, red, n =164 FAs, 5 cells) and αV-GFP-less-constrained (bottom, green n =232 FAs, 11 cells) relative to the FA long axis measured using the Instantaneous FluoPolScope. Circular Gaussian fits to the orientation histogram are outlined in red and green for each chimera, respectively. The reference frame for the long axis of the FAs is the 0-180° axis with the leading edge of the cell towards right. **(g)** (Top) Orientation of the projection (onto the X-Y image plane of the microscope) of the GFP dipole orientations with respect to the αvβ3 integrin headpiece in the Rosetta ensemble of model structures on a unit sphere (each gray point= one Rosetta model) when the integrin headpiece is tilted at ϕ = 12^o^. Microscope frame of reference defined as in **(e).**Lines between the points and the origin give the dipole orientation and their length is proportional to their contribution to polarization factor. Gray, yellow and red spheres represent the points defining the integrin/microscope frame-of-reference as in **(e).**Dashed red and green outlines highlight circular Gaussian fits of experimental data as described in **(f)** for αV-GFP-constrained (red) and αV-GFP-less-constrained (green). Solid red and green outlines highlight the angular range of the circular Gaussian defined by its angular full-width-at-half-maximum (FWHM) used to evaluate the overlap of the Rosetta models with the experimental measurements (Supp. methods). (Bottom) Plot of the fraction of Rosetta models of dipole orientations relative to the (αvβ3 integrin headpiece that lie within the FWHM of circular Gaussian of the experimentally measured distribution of dipole orientations w.r.t the FA long axis as a function of tilt of the integrin headpiece (ϕ, in degrees) when Θ = 0^o^ for αv-GFP-constrained (red) or αv-GFP-less-constrained (green), and their sum (black). Value of ϕ at peak overlap highlighted with a dotted line. **(h)** Speculative model of the alignment of integrins in FA (left, center, view along the microscope Z axis) and the proposed orientations of αVβ3 integrin, talin and actin in FA (right, view along the microscope Y axis) in the frame of reference defined in **(e).**Talin is depicted as a monomer rather than a dimer for clarity.

To relate the measured GFP dipole orientations in FAs for our two chimeras to integrin orientation in FAs, we created an integrin/microscope frame of reference (Fig 4e). The X,Y plane was defined as parallel to the substrate/cell plasma membrane, with the FA long axis aligned along the X axis with +X pointing towards the cell leading edge. The X,Z plane was defined by three conserved positions in the integrin headpiece: one in Asp of the RGD ligand at the origin; and one at each junction between the headpiece and the αV and β3 legs. The ligand-binding site pointed towards the substrate in the –Z direction along the optical (Z) axis.

To model the orientations of GFP relative to αVβ3 integrin for both chimeras, we employed Rosetta^26^. Integrin and GFP were treated as rigid bodies, and alternative conformations for the connecting segments at the integrin-GFP junction that produced no steric clashes or strain in the fusion region were sampled. Rosetta yielded an ensemble of model structures of the two fusion proteins (Fig 4b) that provide a range of orientations in which the actual orientation should be included. This predicted a broader range of possible dipole orientations relative to the integrin for αV-GFP-less-constrained compared to that for αV-GFP-constrained, as expected from the longer fusion junction segments (Fig S6a).

We then modeled the effect of rotating the ensemble of Rosetta structures around the integrinligand point (at the origin) on the orientation of the GFP dipole in the image plane to determine when it matched the experimentally determined dipole orientations in FA for both integrin-GFP chimeras (see supp methods, Fig S6b,c, Supp Movie 1). We assumed *ab initio* that the integrin is aligned with the line defined by the α-leg junction to β-lleg junction along the FA (X) axis, as would be expected if tension were applied to the β-leg via retrograde flow and as confirmed in a companion manuscript (Nordenfeldt et al., submitted under separate cover). We determined the tilt of the integrin headpiece relative to the membrane plane (rotation around Y axis) that maximizes the overlap between the dipole orientations in the Rosetta models and the experimentally measured distribution of dipole orientations for both constructs. This predicted a headpiece tilt of ~0-15° relative to the membrane plane (Fig 4g).

Considering our assumptions and the possible caveats of Rosetta modeling, we thus speculate that αVβ3 integrin headpiece is tilted relative to the membrane-normal (optical) axis along the axis of actin retrograde flow (Fig 4h). This suggests that when bound to ligand and under tension, integrins may be oriented more parallel than normal to the membrane. Remarkably, the ~0-15° angle of integrin relative to the membrane is similar to the tilt of both actin bundles and extended talin molecules in FAs in the leading edge of fibroblasts^27–29^. Whether the alignment of the actin/talin and integrin-ligand systems along a similar angle occur because the normal forces inside the cell are identical or are small relative to the parallel forces^29^ will be an important question for further study.

In conclusion, our study shows that FA and NA are built of an anisotropic molecular scaffold that is actively imposed by the directional forces of F-actin retrograde flow. In the absence of ligation or linkage to the actin filaments, integrin molecules do not exhibit any co-alignment with their neighbors. It is likely that binding to ECM ligands by integrin that attach to oriented, flowing actin filaments in NA and FA^3^, and this in turn generates an ordered molecular ensemble. This actively driven molecular ordering may mediate the ability of integrins to act as directional sensors of force and ECM topology to guide cell migration and mediate proper ECM remodeling during development and wound healing.

## ϕ Acknowledgements

We thank Nikon Instruments and Andor Technology for use of imaging equipment, and Subhasri Ghosh, Bram Prevo, Peter Gross, Weiwei Lou, Daria Bonazzi, Thomas van Zanten, Kabir Husain, Amy Gladfelter, Cheng-Han Yu and Greg Alushin for discussion. Supported by the Lillie Research award from Marine Biological Laboratory and the University of Chicago (CMW, TAS, SM, TT), NIH 5R13GM085967 grant to the Physiology Course at Marine Biological Laboratory, HHMI Summer Institute at Marine Biological Laboratory (SM), NIH CA31798 (TAS, PN, TJM), NIH GM100160 (TT, SBM), NIH GM092802 (DB, KN), NIH GM114274 (RO, SBM), HFSP postdoctoral fellowship LT000096/2011-C (SBM), National Center for Biological Sciences-Tata Institute of Fundamental Research (SM, JMK), HFSP RGP0027/2012, JC Bose Fellowship, and Wellcome Trust DBT-Alliance Margadarshi Fellowship (SM) (SM), and NHLBI Division of Intramural Research (CMW, VS).

## Author Contributions

This project was initiated in the Physiology Course at the Marine Biology Lab in Woods Hole, MA. CMW, SM, TAS, VS, PN, and JMK conceived of the experimental plan. CMW, TAS, TT, RO and SM supervised the project. SBM, TT, and JKM set up polarization microscopes. VS, JKM, and SBM performed imaging experiments and analyzed data. PN made GFP-integrin constructs and PN and TIM analyzed data. RO advised on analysis. KN, TAS, and DB performed Rosetta modeling. TAS constructed the integrin/microscope frame-of-reference and defined the GFP dipole orientation within Rosetta Models. TAS and SBM analyzed Rosetta data, and SBM performed image simulations. CMW and VS drafted the manuscript. All authors discussed the results and commented on the manuscript.

## Supplemental Materials

**Supplementary Table 1.**
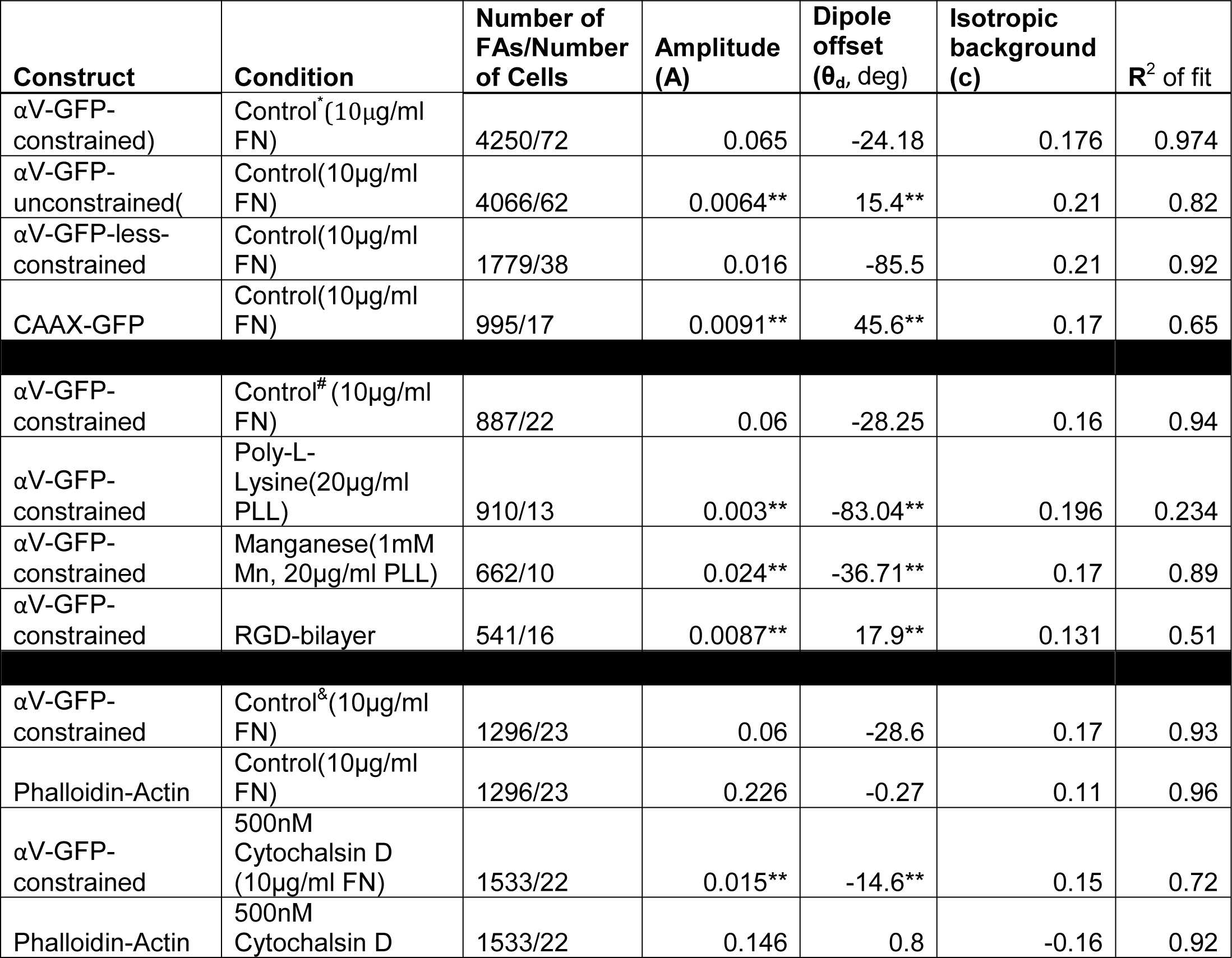
Summary of parameters obtained by fitting anisotropy (r) vs. FA orientation (γ) for data obtained from emission anisotropy total internal reflection fluorescence microscopy (EA-TIRFM). MEF co-expressing GFP-tagged αV integrins, membrane marker, or soluble markers, together with untagged β3 integrins and/or mApple-Paxillin were imaged by EA-TIRFM. Anisotropy magnitude maps were computed from EA-TIRFM images, FA were segmented in the paxillin channel (unless noted otherwise) and the orientation of the FA long axes relative to the axis of excitation polarization determined. Average anisotropy (r) in each FA was plotted against the orientation of that FA (γ) and data was fitted to the equation: ***r= A.cos^2^(γ+θ_d_)+C***. Here, r is the measured anisotropy, A is amplitude of the cosine^2^ function which directly relates to the magnitude of angular dependence of *r* w.r.t polarized light, *γ* is the angle of the long axis of the FA w.r.t to polarized light, *θ*_d_is the angular offset from 0° and C is the isotropic background. *A*, *θ*_d_ and *C* are obtained from the fit (Matlab^®^ curve fitting tool). R^2^ of each fit is also tabulated below.

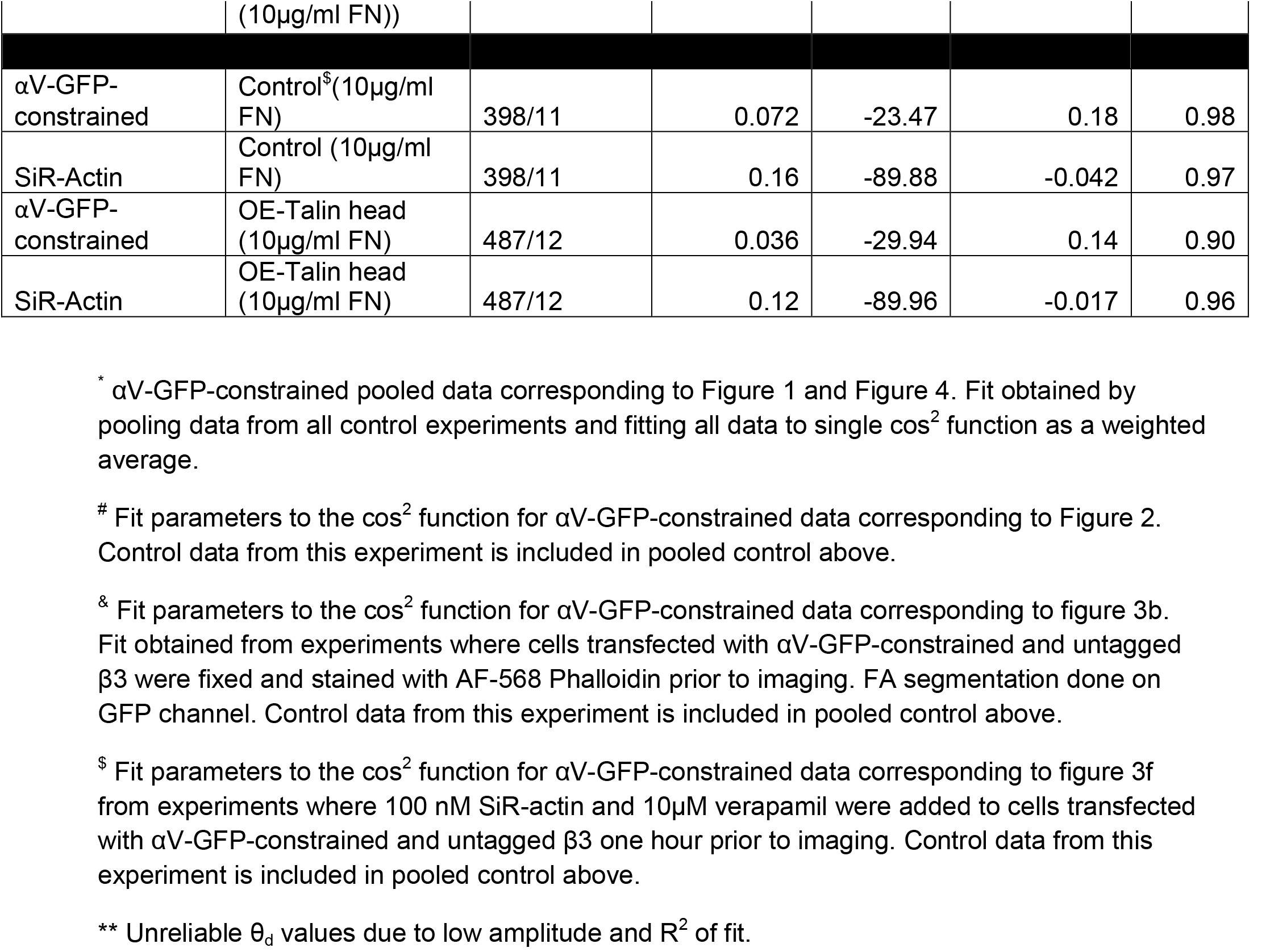

### Supplementary Figure legends

**Figure S1:**
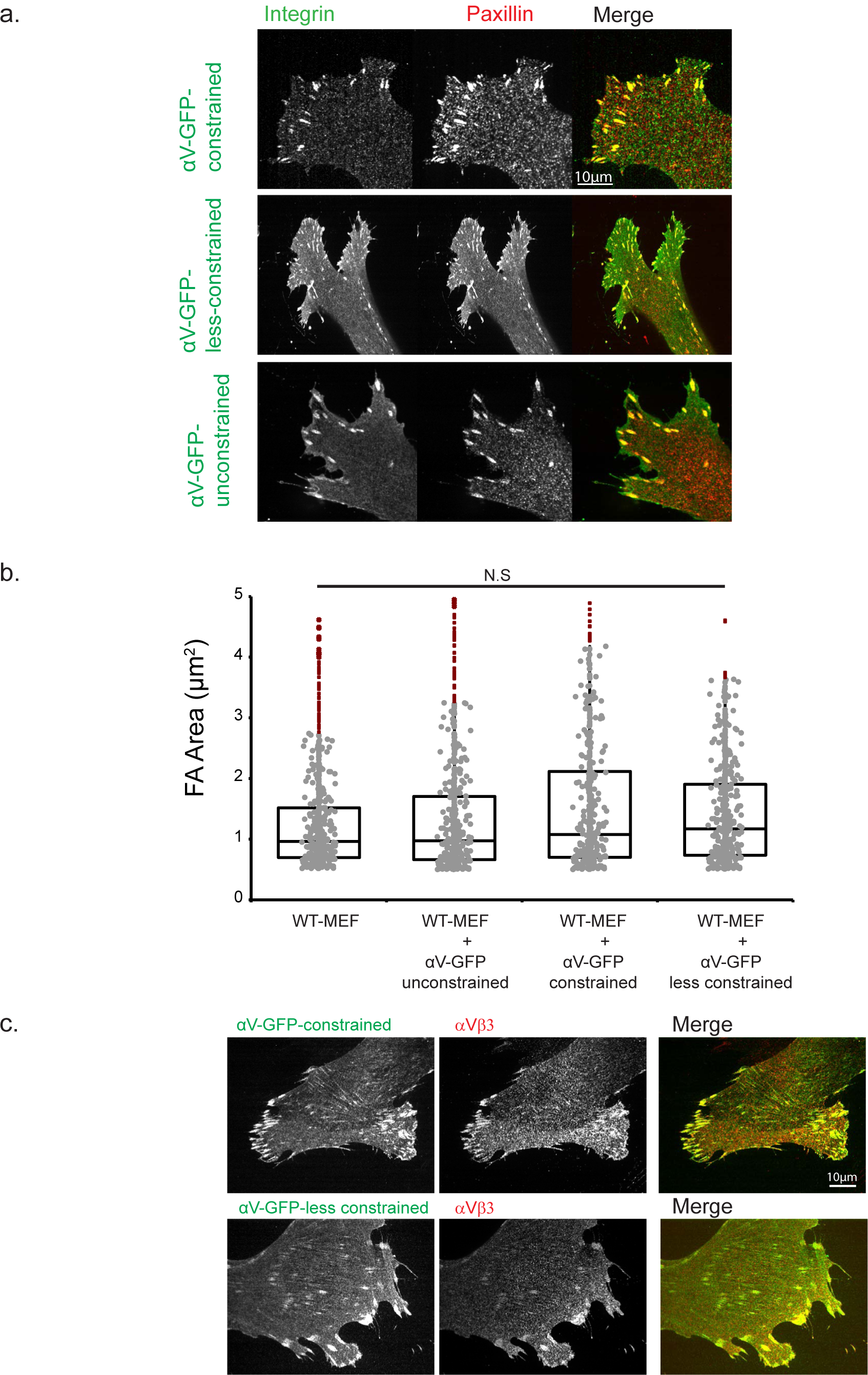
a. **Fluorescently tagged integrin chimeras co-localize with paxillin in FAs.**Representative confocal micrograph of fixed MEF cells expressing either αV-GFP-constrained **(top row, left),** αV-GFP-less-constrained **(middle row, left)** or αV-GFP-unconstrained **(bottom row, left)** together with untagged β3 and immunofluorescence of paxillin **(middle column)**. Cells were transfected and allowed to recover overnight and then were plated on 10µg/ml FN coated coverslips for 4-6 hours prior to fixation for immunostaining. Color merge shown in right column, expressed integrin channel in green and paxillin in red. b. **Expression of integrin chimeras does not affect FA morphology.** Box plot quantification of focal adhesion size (area, um^2^) from analysis of fluorescence images in cells transfected with the indicated αV-GFP-integrin construct, co-transfected with untagged β3. FA area was obtained by thresholding the paxillin channel. Box plots: gray points, individual FAs: red points, outliers; box, 95% confidence interval; bar, median. N= 500-700 adhesions. NS- Not significant. Kruskal-Wallis H test. c. **Expressed αV integrin-GFP chimeras conjugate with β3 subunit**. Representative confocal micrograph of fixed MEF cells expressing either αV-GFP-constrained **(top row, left, green)** or αV-GFP-less-constrained **(bottom row, left, green)** together with untagged β3 and immunofluorescence of αVβ3 integrin using LM609 antibody **(top and bottom row, middle panel, red)**. Cells were transfected with the indicated construct and allowed to recover overnight before plating on coverslips coated with 10µg/ml FN for 4-6 hours prior to fixation. Color merge shown in right column. Scale bar= 10µm.

**Figure S2:**
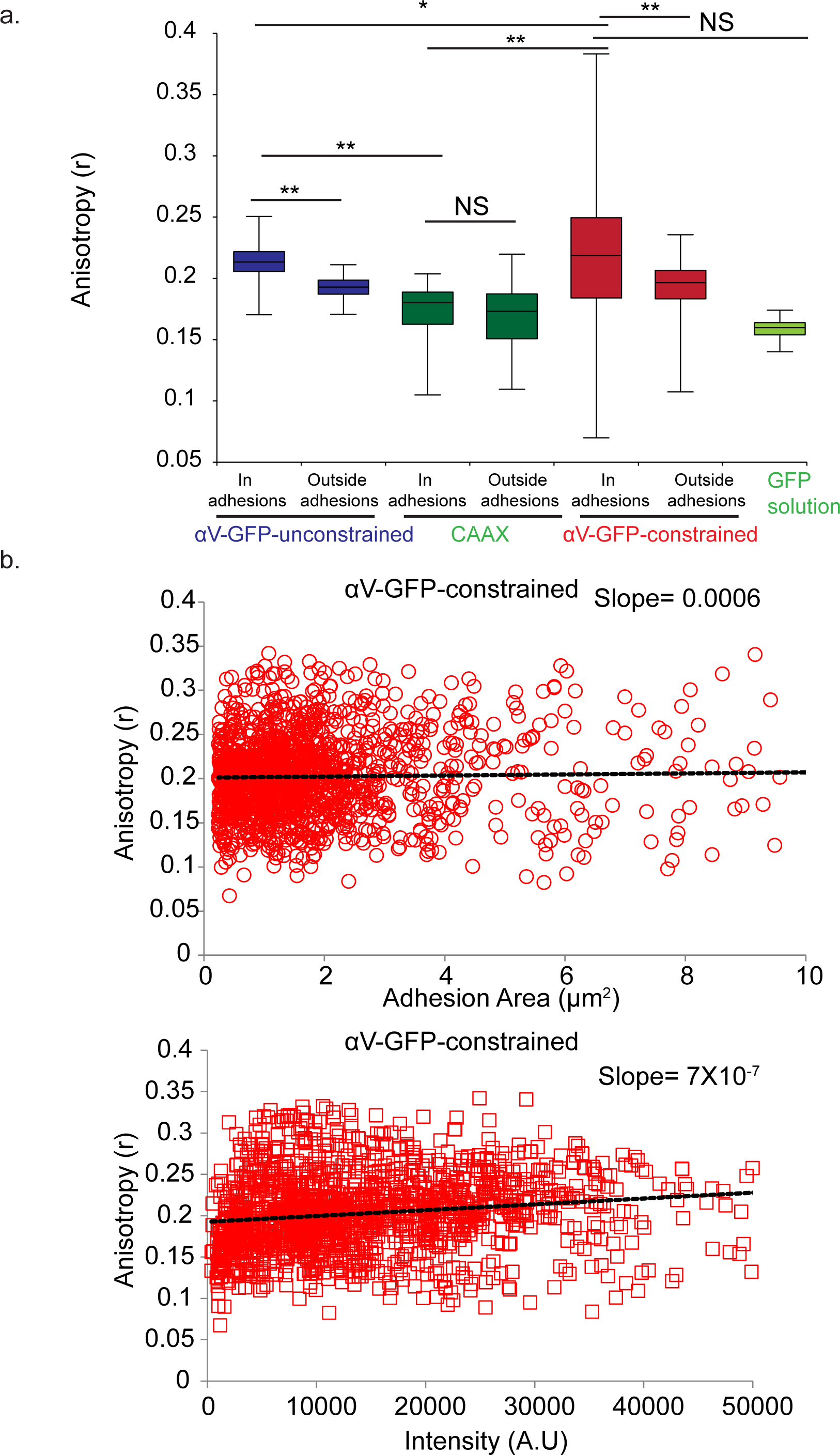
(a) **Integrin emission anistropy is higher within FAs than cellular areas outside FAs**. Box plot of average emission anistropy of expressed αV-unconstrained (blue), EGFP-CAAX (green) or αV-GFP-constrained (red) in segmented FAs or ROIs outside FAs and emission anistropy of GFP solution (light green). Both αV-GFP integrin chimeras were co-transfected with untagged β3 and mApple-tagged Paxillin. Cells expressing EGFP-CAAX were co-transfected with mApple-tagged Paxillin. Cells were then subjected to EA-TIRFM imaging. For all conditions, the mApple-tagged paxillin channel was used to segment FAs. Box plots: gray points, individual FAs: red points, outliers; box, 95% confidence interval; bar, median. N= 900-1500 adhesions. **-p<0.0001, *-p<0.01, NS- Not significant. Kruskal Wallis H test. (b) **Emission anistropy of αV-GFP-constrained is independent of FA size**. MEF co-expressing αV-GFP-constrained, untagged β3 and mApple-tagged Paxillin were subjected to EA-TIRFM imaging and emission anistropy maps computed. FAs were segmented using the mApple-paxillin channel and used to find the average emission anistropy of αV-GFP-constrained and area of each FA. Plot of average FA emission anisotropy (r) of αV-GFP-constrained versus FA area of the FA. Linear fit of data shown in black dashed line. (c) **Emission anistropy of αV-GFP-constrained is independent of average intensity/signal at FAs.** Plot of average FA emission anisotropy (r) versus average FA intensity for αV-GFP-constrained. MEF co-expressing αV-GFP-constrained, untagged β3 and mApple-tagged Paxillin were subjected to EA-TIRFM imaging and emission anistropy maps computed. FAs were segmented using the mApple-paxillin channel and used to find the average emission anistropy of αV-GFP-constrained in each FA. The average intensity of αV-GFP-constrained signal ((I_pa_+I_pe_)/2) in FAs was computed and recorded along with emission anistropy. Linear fit of data shown in black dashed line.

**Figure S3.**
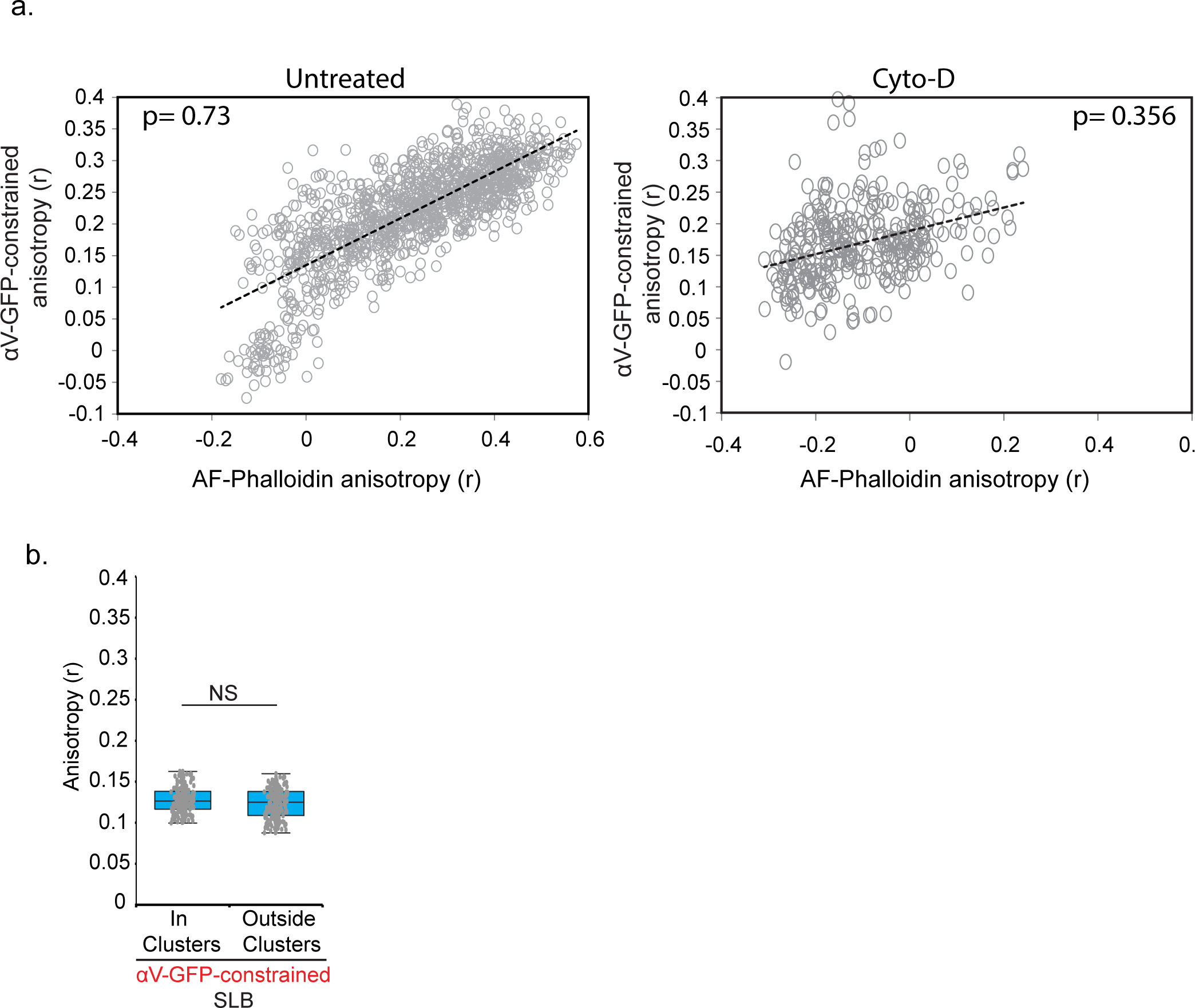
(a) **Emission anistropy of αV-GFP-constrained correlates with emission anistropy of F- actin in FAs**. Plot of αV-GFP-constrained emission anistropy vs AlexaFluor561-Phalloidin emission anistropy in FAs in **(left)** untreated condition or **(right)** treated with 500nM Cytochalasin-D. Cells were transfected with αV-GFP-constrained, co-transfected with untagged β3 and allowed to recover overnight after transfections. Cells were then plated on 10µg/ml FN coated coverslips for 4 hours after which cells were either treated with 500nM Cytochalasin-D and then fixed **(right)** or directly fixed and stained with AF-561 phalloidin **(left)**. Cells were then subjected to EATIRFM imaging and emission anistropy maps computed. FAs were segmented using the αV-GFP-constrained channel and used to find emission anistropy of αV-GFP-constrained and phalloidin-labeled actin in each FA and plotted. Linear fit of data shown in black dashed line. p: co-efficient of correlation. (b) **Mobile ligand on SLB reduces integrin emission anistropy in** αV-GFP-constrained **clusters to background level.** Cells were transfected with αV-GFP-constrained, untagged β3 and mApple-tagged Paxillin and allowed to recover overnight. Prior to the experiment, cells were trypsinized and plated on supported lipid bilayers (SLB) as described in methods and subjected to EA-TIRFM imaging 30 minutes after plating. Clusters were segmented using the paxillin channel. Box plot of average emission anistropy(r) of αV-GFP-constrained in clusters or in ROIs outside clusters. Box plots: gray points, individual FAs, box, 95% confidence interval; bar, median, error bars, standard deviation. N= 400-450 clusters, NS- Not significant, Mann Whitney U test.

**Figure S4.**
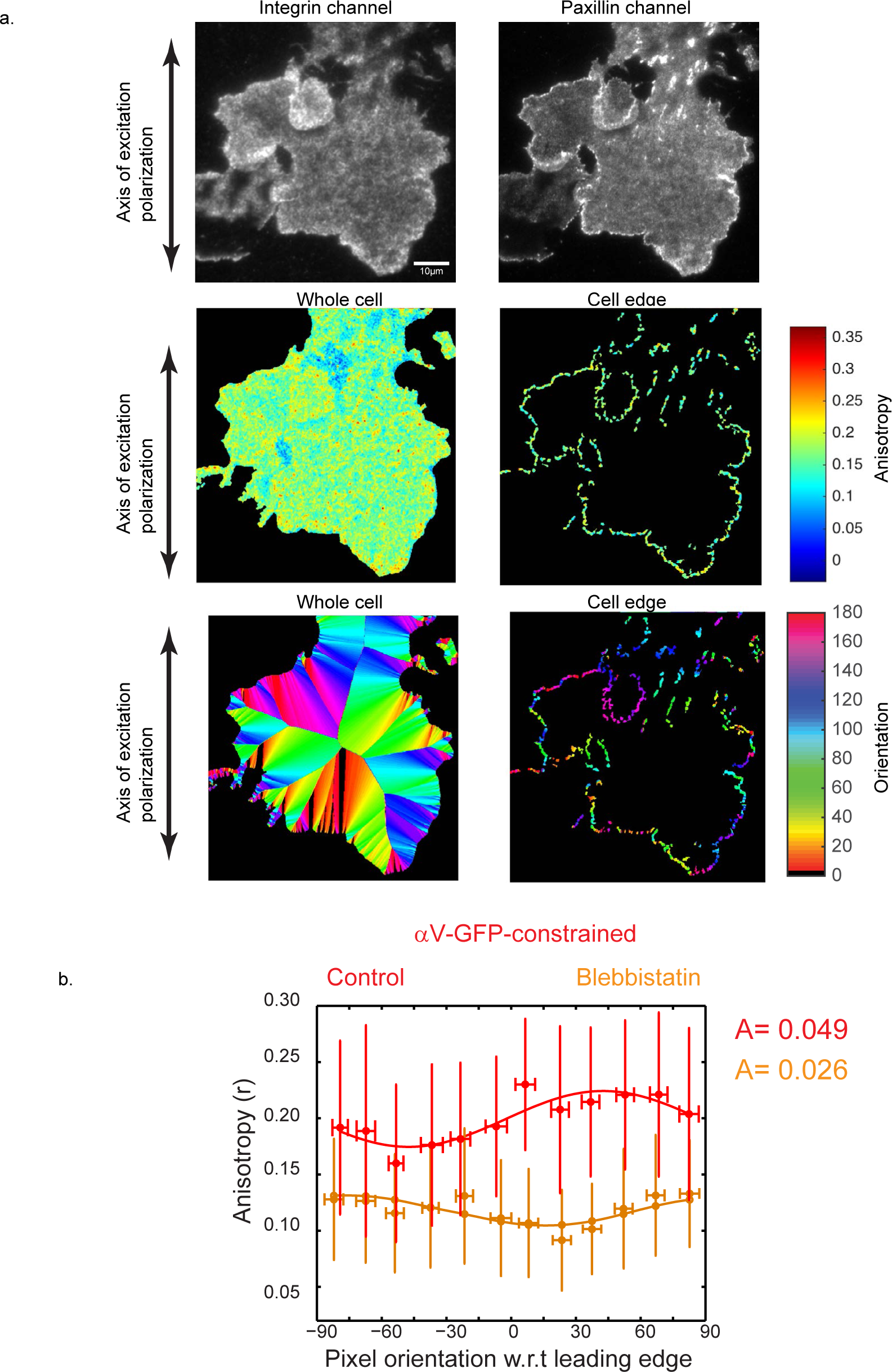
**a. Analysis pipeline of αV-GFP-constrained emission anistropy relative to the orientation of the nearest leading edge in blebbistatin treated cells. (top)** Fluorescence micrographs of cells expressing αV-GFP-constrained **(left)** and mApple-Paxillin **(right)**, co-transfected with untagged β3 and treated with 50µM Blebbistatin for one hour, then subjected to EA-TIRFM imaging.**(Middle row, left)** Emission anistropy map of the segmented area of the whole cell. **(Middle row, right)** Emission anistropy map of the cell edge generated by eroding the whole cell mask by 10 pixels and then inversely combining with the whole cell mask in (middle, left). **(Bottom row, left)** Orientation assignment of all pixels in a cell. Each pixel in a cell is an assigned an absolute orientation for the vector normal to the closest cell edge. **(Bottom row, center)** Orientation assignment of pixels at the cell edge generated by using the cell edge mask in (middle row, right). Scale bar= 10µm.**b. Angular dependence of the emission anistropy of αV-GFP-constrained as a function of leading edge orientation in blebbistatin treated cells**. (orange) Emission anistropy(r) value for αV-GFP-constrained at pixels at the cell edge (a, middle, right) plotted against the orientation assignment of the pixel relative to the cell edge (a, bottom, right) for the cell shown in **(a).** Similar analysis of a cell in untreated condition shown in red. Pixel orientation was binned at 15° increments. Note that because the anisotropy values per pixel data is plotted relative to the orientation of the nearest leading edge, as opposed to all other experiments where it is poltted relative to the axis of excitation polarization), no values of θ_d_ are reported.

**Figure S5.**
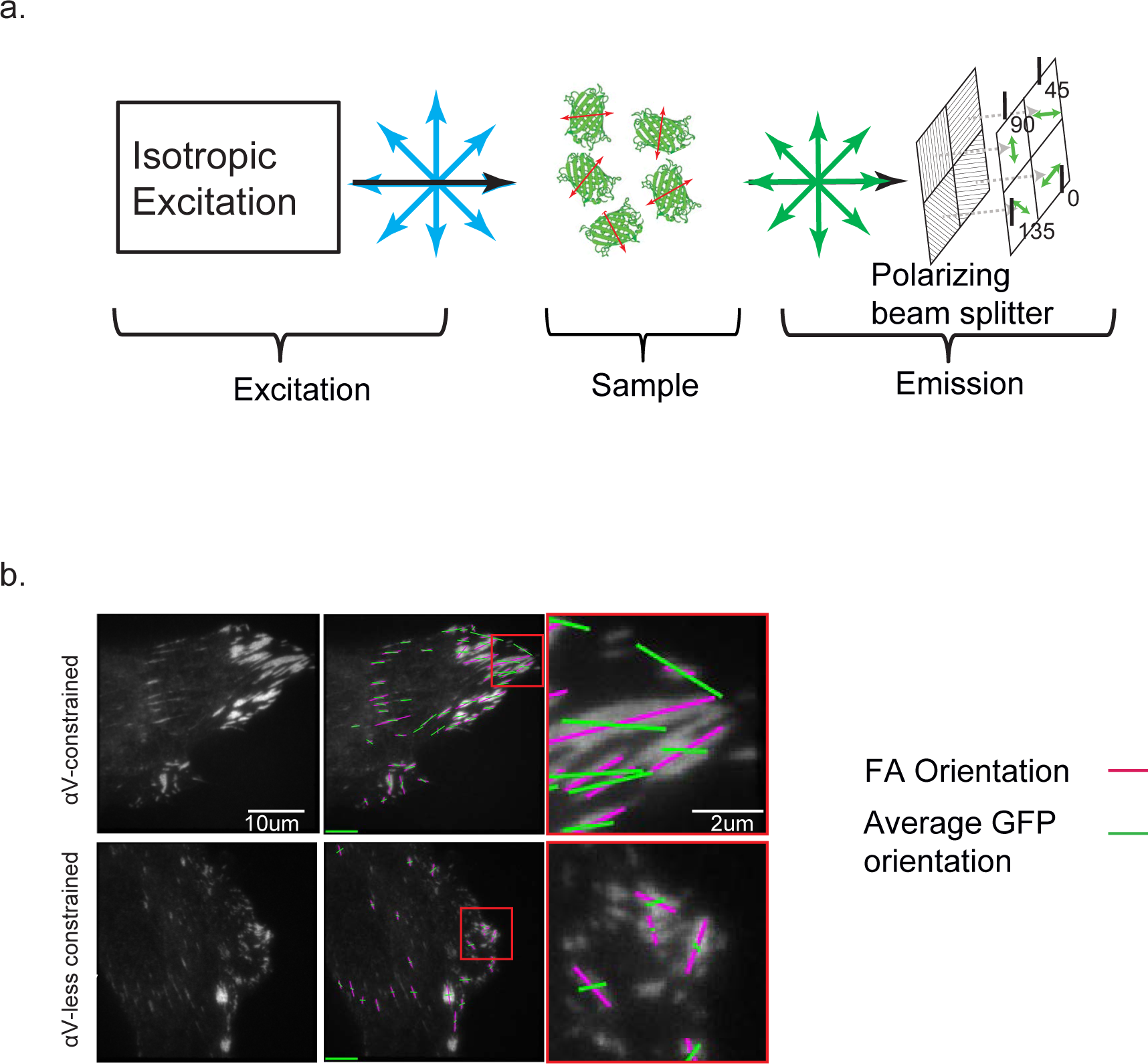
a. **Schematic of Instantaneous FluoPolScope setup.** b. **Direct measurement of orientation of the GFP dipole for αV-GFP-constrained and αV-GFP-less-constrained using Instantaneous FluoPolScope.** Cells were co-transfected with αV-GFP chimeras, untagged β3 and mCherry paxillin and subjected to imaging on the instantaneous FluoPolScope. Representative fluorescence micrograph of αV-GFP-constrained **(top, -left)** or αV-GFP-less-constrained **(bottom, left). (Top -right, bottom -right)** Long axis of the FA (magenta, determined from segmentation in the paxillin channel) and the computed average orientation of GFP dipoles (green). Scale bar= 10µm.**(top right, bottom right)** Zoom in from panels to the immediate left.

**Figure S6.**
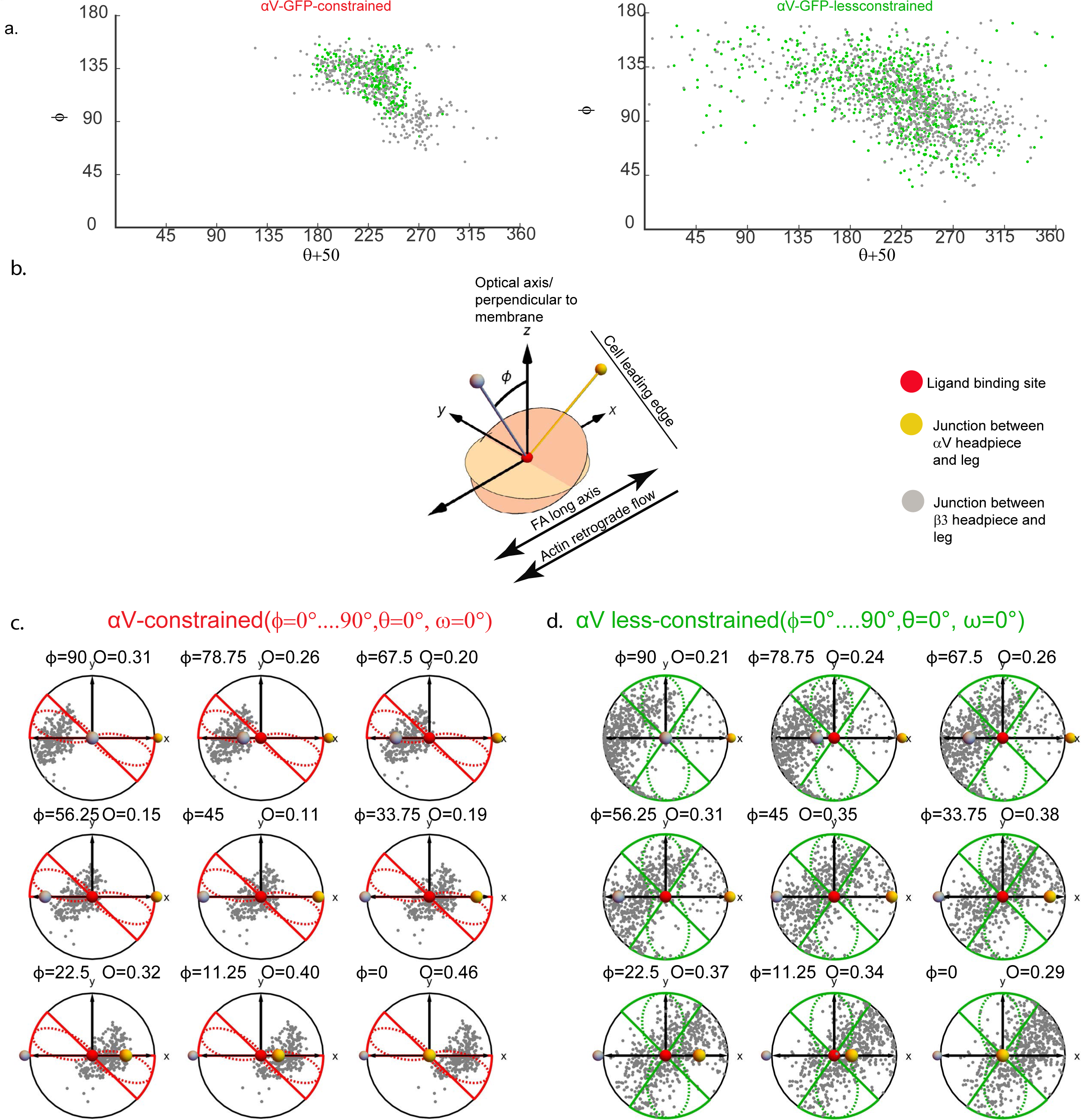
**a. Determination of the orientation of the GFP dipole relative to the integrin frame of reference for the ensemble of Rosetta models of GFP-integrin chimeras.** For each model structure of the integrin-GFP chimera that was output by Rosetta, the orientation of the GFP dipole was determined relative to the integrin headpiece. The dipole was defined by a vector from the N atom of Val112 to the O atom of Ser147. A plane in the integrin frame-of-reference was defined by 3 Cα atoms: Arg4380 at the junction between the headpiece and the αV leg, Pro111 at the junction between the headpiece and the β3 leg, and Asp of the RGD ligand. The plane spanned by the 3 Cα atoms was made the X-Z plane, the ligand Cα atom made the origin of the integrin frame-of-reference, and the line from the ligand point to the leg junction was made the X-axis. The Z-axis is then almost parallel to the line between the ligand point and the β leg junction (see Methods for exact coordinates). The orientation of the dipole vector relative to this plane was denoted as (ϕ,θ), where ϕϕis the tilt relative to Z-axis and θ is the azimuth in the XY plane of the integrin/microscope frame-of-reference. Each point in the graphs represents this orientation of the GFP dipole from one Rosetta model. The Rosetta models were ranked according to the energy and models with energies less than 40 percentile are shown by GREEN dots while the rest of the models are shown by GRAY dots. (red, left) αV-GFP-constrained, (right, green) αVGFP-less-constrained. **b. Schematic of the coordinate system for tilting the GFP-integrin chimeras in the microscope frame-of-reference along the retrograde flow/FA axis.** Coordinates defining the integrin/microscope frame-of-reference are shown as yellow (headpiece-/ leg junction), grey (headpiece-β leg junction) and red (ligand) circles. At the start of the simulation in b, the integrin headpiece was aligned with the frame of reference (XYZ) as follows: The cell leading edge was to the right (+x), the focal adhesion long axis was along the x axis, the ligand (red circle/sphere) was held at the origin and towards the substrate (-z) and the legs oriented at (+z), the α leg was anterograde (+x) and the β leg retrograde (-x), because flow is assumed to pull the β leg retrograde. The tilt of the integrin-GFP chimeras in the microscope frame-of-reference towards the retrograde flow gives rise to angular rotation (*ϕ*) around the y axis. **c, d. Simulations of the effect of specific tilts of αVβ3 integrin along the retrograde flow/FA axis on the overlap of dipole orientations of Rosetta models and experimentally measured dipole distribution for αVβ3 integrin-GFP chimeras.** (c, red) αV-GFP-constrained. ( **d,** green) αV-GFP-less-constrained. In the plots, each gray point represents GFP dipole orientation in the microscope frame of reference for one Rosetta model, such that its position around the circumference represents its angle with respect to the FA long axis, and its distance from origin represents its tilt around the y axis in the direction of retrograde flow (-x) (*ϕ* angle as defined in (b)). Each plot shows the projections of 3D dipole orientations at a specific tilt *ϕ* (denoted above each plot) of the integrin-GFP headpiece. Simulations were performed at increments of *ϕ* = 11.5^o^. The circular Gaussian fits to the experimentally measured GFP dipole orientations are overlaid by a dashed line. The angular range of the circular Gaussian is defined by its full-width-at-half-max (FWHM) and overlaid by a solid line. The overlap *O* (denoted above each plot) between the angular range of Rosetta models and the angular range of experimentally determined GFP orientation is computed as described in Methods.

### Supplemental Video Legend

#### Video 1

Simulation (Corresponding to **Figure 4(g)** and **Figure S6**) of the 2D projection (onto the X-Y image plane of the microscope) of the predicted GFP dipole orientations in Rosetta model structures on a unit sphere (each gray point= one Rosetta model) when the integrin headpiece is tilted from *ϕ* = 90^o^(No tilt) to *ϕ* = 0^o^(high tilt). Microscope frame of reference defined as in **Figure 4 (e).** Gray, yellow and red circles represent the points defining the integrin plane-of-reference as described in the integrin headpiece. Dashed red and green outlines highlight circular Gaussian fits of experimental data from Instantaneous FluoPolScope for αV-GFP-constrained (Left, red) and αV-GFP-less-constrained (Right, green). Solid red and green outlines highlight angular range (full-width-at-half-maximum or FWHM) of the circular Gaussian. Top view corresponds to top-down microscope view along the optical axis. Bottom corresponds to perspective view. Overlap parameter as described in the Methods is shown.

### Materials and Methods

#### cDNA expression vectors

For the integrin-GFP fusion constructs, the insert position of GFP was based on previous work^1^ where cutinase was attached in frame to αV. All integrin-GFP constructs were made using three-segment (A,B,C) overlap PCR with wild type human αV cDNA and either pEGFP-N1 (Clontech, for for **αV-GFP-less-constrained**) or moxGFP^2^ (for **αV-GFP-constrained**) as sources for GFP cDNA. After the three segments had been made and joined together using PCR (Accuprime Pfx, high-fidelity polymerase, ThermoFisher), the complete A–C sequence and the wild type αV-pcDNA3.1 plasmid were cut with restriction enzymes (New England Biolabs) and ligated together with T4 ligase (Roche) after de-phosphorylation (rAPID alkaline phosphatase, Roche) and purification (Qiagen) of the linearized plasmid. The overall plasmid integrities were verified with size matching of multi-site single restriction enzyme digestion compared to virtual digest patterns (Serial Cloner) and the inserts were verified by full sequencing. Surface expression of the αV-GFP constructs were validated by transient co-expression with β3 in 293T cells. The base design was full-length EGFP inserted between Lys-259 and Asn-260 in the αV β propeller domain, with an addition of QG and GG to extend the loop out from the β propeller with residues from the analogous loop in α_L_. This is referred to as **αV-GFP-constrained** and the primers used were:

*A1: 5′- AGA TGT GGT TCT AGA GCC ACC ATG GCT TTT CCG CCG C -3′; A2: 5′- TCG CCC TTG CTC ACC ATG CCC TGC TTC CCA TCA TAA ATA TAA ACC ATT CCC -3′; B1: 5′- TAT ATT TAT GAT GGG AAG CAG GGC ATG GTG AGC AAG GGC GAG -3′; B2: 5′- TAT AAG GAG GAC ATG TTT CCT CCC TTG TAC AGC TCG TCC ATG CC -3′; C1: 5′- ATG GAC GAG CTG TAC AAG GGA GGA AAC ATG TCC TCC TTA TAC AAT TTT ACT GG -3′; C2: 5′- ACT CTT AGT AGC GGC CGC TTA AGT TTC TGA GTT TCC TTC ACC ATT TTC -3′.*

By increasing the linker length, we made the less constrained chimera (**αV-GFP-less constrained)** and the primers used were:

*A1: 5′-AGA TGT GGT TCT AGA GCC ACC ATG GCT TTT CCG CCG C -3′; A2: 5′- TCA CCA TGC CAG ATC CAG AGC CCT GCT TCC CAT CAT AAA TAT AAA CCA TTC CC -3′; B1: 5′- TGA TGG GAA GCA GGG CTC TGG ATC TGG CAT GGT GAG CAA GGG CGA G -3′; B2: 5′- AGG ACA TGT TTC CTC CGC TGC CTG AGC CCT TGT ACA GCT CGT CCA TGC C -3′; C1: 5′- AAG GGC TCA GGC AGC GGA GGA AAC ATG TCC TCC TTA TAC AAT TTT ACT GG -3′; C2: 5′- ACT CTT AGT AGC GGC CGC TTA AGT TTC TGA GTT TCC TTC ACC ATT TTC -3′.*

cDNAs encoding eGFP- αV integrin(**αV-unconstrained**), mApple paxillin, eGFP-Actin, EGFPCAAX and soluble GFP were the kind gift of M. Davidson (Florida State University, Tallahassee, FL) and have been described previously^2,3^. Untagged β3 integrin was provided by Mark Ginsberg (UCSD, San Diego, CA).

#### Cell culture and imaging sample preparation

Mouse embryonic fibroblasts (MEFs) were maintained in DMEM supplemented with 10% FBS (Invitrogen), 2mM glutamine, and 100 units/ml of penicillin/streptomycin (Invitrogen) at 5% CO_2_ as previously described^4^. For experiments, cells were transfected by nucleofection with endotoxin-free expression vector DNA (~1µg per 10^6^ cells) per manufacturer’s protocol (Lonza). Cells transfected with αV integrin constructs were co-transfected with untagged β3 to promote co-translational dimerization. Transfected cells were cultured overnight and re-plated on Mat Tek petri dishes (Mat Tek Corp.) that had been coated with 10µg/ml fibronectin (Millipore Corp., unless coating is specified otherwise) 4 hours prior to imaging. Cells were imaged live in imaging media (phenol red-free DMEM supplemented with 10% FBS (Invitrogen), 2mM glutamine, 100 units/ml of penicillin/streptomycin, 20 mM Hepes (Invitrogen) and 10 U/ml Oxyrase (Oxyrase Inc). For experiments with phalloidin staining, cells were fixed with 4% paraformaldehyde in cytoskeleton buffer (CB, 10mM MES pH 6.1, 138mM KCl, 3mM MgCl, 2mM EGTA) for 30 minutes at 37°C and then permeabilized for 5 min with 0.25% Triton X-100 in CB. Free aldehydes were reacted with 10mM glycine and then washed in TBS. Cells were then incubated with fluorophore-conjugated Alexa fluor 568 phalloidin (Thermofischer Scientific) for 1 hour and then washed and imaged in TBS. For cytochalasin-D experiments, cells plated for 4 hours on Mat Tek dishes were treated with 500nM cytochalasin-D (Sigma-Aldrich) for 15-30 minutes before fixing and staining with phalloidin as described above. Blebbistatin experiments were done similarly with cells treated with 50µM blebbistatin (Tocris Biosciences) for 1 hour before fixation and imaging. Poly-L-Lysine (PLL) experiments were done by coating Mat Tek dishes with 20µg/ml PLL (Sigma Aldrich) for 1h and then plating cells for 1 hour before imaging. For Mn^2+^ experiments, cells were plated on PLL coated Mat Tek dishes with 1mM MnCl_2_ for 2 hours before imaging. The SiR actin kit was obtained from Cytoskeleton Inc. and used according to manufacturer’s protocol. Briefly, cells expressing αV-constrained construct and mApple-paxillin were plated for 4 hours on FN coated Mat Tek dishes and then incubated with 100nM SiR-actin and 10µM verapamil diluted in regular growth media for 1 hour. Prior to imaging, the media was replaced with imaging media and imaged immediately. Supported lipid bilayers were generated as previously described^5^. Briefly, 1,2-dioleoyl-sn-glycero-3-phosphocholine(DOPC) doped with 1,2-dipalmitoy-sn-glycero-3-phosphoethanolamine-N-(cap biotinyl)(16:0 Biotinyl Cap PE, Avanti Polar Lipids Inc.) were used to assemble SLBs and then functionalized with biotin RGD using labeled Dylite neutravidin as a linker. For FSM imaging of actin flow, MEF cells were transfected with EGFP-actin (0.5µg/ml) and mApple-paxillin (1µg/ml) and directly plated on FN (10µg/ml) coated coverslips for 4 hours in complete media. Prior to imaging, coverslips were washed with PBS and imaged in imaging media.

### Image Acquisition

#### Emission Anisotropy Total internal reflection microscopy (EA-TIRFM)

All images were acquired using the Total Internal Reflection Fluorescence Microcopy (TIRF) mode on a Nikon Eclipse TiE inverted microscope equipped with the Perfect Focus System (PFS3;Nikon, USA), a motorized TIRF illuminator (Nikon, USA) and a motorized stage (TI-S-ER Motorized Stage with Encoders; Nikon, USA). Illumination was provided by a multi-wavelength (405 nm[15-25mW], 488nm [45-55mW], 561 nm[45-55 mW], 640 nm[35-45 mW]) polarization-maintaining fiber coupled Monolithic Laser Combiner (Model: MLC400, Agilent Technologies). This arrangement resulted in the generation of a polarized TIRF evanescent field at the sample plane^6^.

Images were collected with a fixed magnification using a 100X Plan Apo 1.49 NA TIRF objective (Nikon, USA) and a 1.5X tube lens to yield a final pixel size corresponding to 109nm on the cameras described below. The typical TIRF illumination depth using the 488 nm was 150-200nm.The emission filters (ET525/50, ET600/50 and ET700/75 band-pass emission filters; Chroma Technology Corp, USA) were mounted onto a motorized turret below the dichroic mirror(405/488/561/638 TIRF Quad cube; Chroma Technology Corp, USA).

The fluorophores were excited with a polarized evanescent TIRF field and the polarized emission was split into the constituent *p* and *s*-polarized components using a high performance nano-wire grid polarizing beam splitter (TR-EMFS-F03; Moxtek Inc.,USA). The resulting I *pa*(parallel) and I *pe*(perpendicular) images were captured on separate, orthogonally placed, iXon Ultra 897 EMCCD cameras (Andor Technology, Belfast, Northern Ireland) using the TuCam two-camera imaging adapter (C-Mount Version [S-CMT]; 1X Magnification [TR-DCIS-100]; Andor Technology, Belfast, Northern Ireland). Images were acquired on the EM-CCDs using the Nikon Imaging Software Elements (NIS Elements Advanced Research; Nikon,USA) with a dual-camera plug-in using a 1 second integration time taken in the electron multiplying gain mode.

**Instantaneous Polscope** *(Manuscript under Review Describing this system is attached)*

A custom microscopy platform using mechanics from Newport Corp was built on an optical table. Laser beams (Coherent Sapphire 488nm and Melles-Griot 561nm) were routed through custom optics and focused on the back focal plane of a 100x 1.49 NA objective (Nikon 100x ApoTIRF 1.49NA). The objective was placed on a Piezo Z-collar (PI P-721 PIFOC) for precise focusing. The laser beams were circularly polarized using a combination of a half wave plate and a quarter wave plate. To achieve isotropic excitation within the focal plane and along the optical axis of the microscope, the circularly polarized laser beam was rapidly rotated (300-400Hz) in the back focal plane of the objective with a large enough radius to achieve total internal reflection at the specimen plane. A dual-band dichroic mirror (Semrock Di01-R488/561) was used to separate laser lines (reflected) and emissions corresponding to GFP and mCherry (transmitted). The specific emission channel was selected using bandpass filters mounted in a filter wheel (Finger Lakes instruments). A quadrant imaging system as described in (attached related manuscript, Mehta et al) was used for instantaneous analysis of fluorescence emission along four polarization orientations at 45° increments (I_0_, I_45_, I_90_, I_135_). Dual-channel imaging of live cells was performed using Micro-Manager (version 1.4.15). All images were acquired using an EMCCD camera (Cascade II: 1024; Photometrics, Tuscon, AZ) operated in the 5MHz readout mode using EM gain.

### Fluorescent speckle microscopy (FSM) and image analysis

TIRF microscopy was performed on an inverted microscope system (Eclipse Ti; Nikon) equipped with the Nikon PerfectFocus system, a servo-motor controlled X-Y stage and a PZ-2000 Piezo Z stage (Applied Scientific Instrumentation) using a 100X/1.49 NA Apo TIRF objective lens. Illumination was provided by a 150mW 488 or a 150mW 546nm laser (Coherent) in a custom-designed laser combiner (Spectral applied research, Richmond ON). Illumination was delivered to the TIRF illuminator (Nikon) by a single mode optical fiber (Oz optics). Images were captured every 3-5 seconds. An appropriate multi-bandpass mirror (Chroma Technology Corp., USA) and single bandpass emission filters (Semrock technology) were used to select emission wavelength. The microscope was controlled using Metamorph software (Molecular devices), temperature was maintained at 37°C (airstream incubator: Nevtek), and images were acquired using a cooled charge coupled device (CoolSNAP MYO, Photometrics).

Speckle tracking in time-lapse movies was done using the qFSM software package from the Danuser lab as previously described^7^. F-actin flow vectors, output by flow analysis of speckle tracks, with signal to noise ratio of >2 were kept for further analysis. The mApple-paxillin channel was used to segment FA and the FA orientation was obtained as described below by modifying an image filtering algorithm developed for blob segmentation by the Danuser lab^8^. Mean F-actin flow velocities in segmented FAs was obtained from F-actin flow vectors and overlaid on the segmented FA maps.

### Image Analysis for EA-TIRFM

The background-subtracted I*pa* and I*pe* images (defined above) were aligned using a custom written affine-based automatic transformation routine in Matlab (Mathworks,USA). To correct for the inherent bias in detection of the polarized components of the emission in the optical path, the G-factor was computed from the I*pa* and I*pe* images of an aqueous solution of fluorescein (pH 11) as follows:

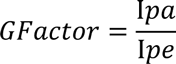

To compute the emission anisotropy of individual FAs, the intensity in the paxillin channel was used to segment the FA using a Matlab code (blobSegmentThreshold.m) from the Danuser Lab. The FA mask was overlaid onto the Ipa and Ipe images in the GFP channel and the corresponding mean intensities were extracted to compute the mean FA anisotropy as follows:

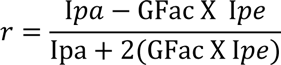

The orientation of each FA was defined by fitting the segmented FA with an ellipse and determining the angle of the ellipse long axis with respect to the plane of polarization excitation (Y axis of the image). The mean emission anistropy and orientation data for individual FAs from all images of all cells were then binned into 15° orientation bins and the mean and standard deviation for the emission anistropy values for each of the bins plotted against the mean orientation as a 2D-scatter plot. The weighted-spline fitting toolbox in Matlab(Mathworks, USA) was used to fit the data to the cosine square equation:

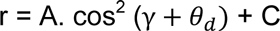

where *r* is the emission anistropy, *A*, the amplitude is a measure of the extent to which fluorophores are aligned within x-y axis of the microscope in the FA, γ is the angle of the FA with respect to the excitation polarization axis, *θ*_*d*_, the angular offset from γ, indicates the angle of the FA long axis relative to excitation polarization axis at which maximal emission anistropy occurs, and *C* is the emission anistropy offset due to background. Angles counter-clockwise from γ were considered as positive angles.

To account for day to day variability, imaging of control samples (αV-GFP-constrained, untreated) were done separately for every perturbation experiment and comparison of perturbation fit parameters was done with those specific controls (Figs 2, 3). Standard deviation reported in the main text corresponds to the average *A* and *θ*_*d*_ across all the different control experiments. Plots for αV-GFP-constrained (in Figure 1 and 4) was obtained by fitting all data across multiple experiments with a single fitting curve, parameters of which are shown in Figure 1e.

### Image analysis of the angular dependence of nascent FA emission anisotropy in blebbistatin-treated cells

Nascent FAs (NA) in cells treated with blebbistatin lose their asymmetric shape and reduce to circular diffraction-limited puncta around the leading edge. This loss of asymmetry prevented FA orientation assignments using the segmentation routine and analysis described above. Thus, based on the assumption that FAs are generally oriented normal to the closest leading edge, instead of determining the emission anisotropy in NA as a function of FA orientation, we determined the emission anisotropy in NAs as a function of the orientation of the closest leading edge. EA-TIRFM image analysis for cells treated with blebbistatin (Figure 3h,3i,3j) was mainly carried out using Matlab 2014a and functions handling all steps were developed in-house. Algorithms are available upon request. Briefly, a background mask was generated by thresholding at 3 standard deviations above background, where the background intensity distribution was estimated by fitting the “left half” of a Gaussian function to the left shoulder of the image intensity histogram. The mask was then used to find and subtract the average background intensity on a frame-by-frame basis on G-factor corrected, aligned images, and the emission anisotropy (r) was determined in each pixel. For emission anisotropy calculations, the data was also pre-filtered with a 3x3 intensity-weighting filter, and to minimize artifacts from division of small integers, only pixels that had a value above four times the background standard deviation in that image frame were used.

For generating a cell edge mask and to assign the orientation of each pixel in the mask relative to the closest cell edge, first the background masking described above was used to initialize cell segmentation independently for each cell. The cell regions were slightly expanded (5-pixel dilation operation) to make sure that background was included. Either active contour segmentation or intensity distribution-based threshold segmentation was used to produce an initial cell mask. Mathematical morphology (a closure operation with a radius of 1 pixel, small object removal, and filling of holes) was applied to further refine these masks, producing accurate cell outlines. For edge segmentation, the cell mask was eroded by 10 pixels, and then inversely combined with the original mask to generate an edge mask. Given the variable cell shapes, an orientation mapping algorithm was devised that would assign relative orientation values in a reproducible manner. It is based on the vector normal to the closest edge in geometric shapes. For circular cells, it yields an orientation axis of 0 to π, that falls along the polarization axis, but for irregular shapes such as polarized cells, the orientation assignment is not comparable across cells, only within. This also means that for non-circular objects the orientation values are not correlated with the polarization axis, and thus the values γ and *θ*_*d*_ for the fit equation described above are arbitrary, and thus not comparable with all non-blebbistatin experiments. However, the value *A*,which is independent of orientation of the cell or polarization axis, is comparable between all experiments and is thus reported in Figure 3. The cell masks were then smoothed with a 3-pixel radius closure operation followed by an Euclidian distance transform of the inverse mask,

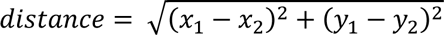

with *x* and *y* pixel coordinates. A numerical 2-dimensional gradient was calculated from the distance transform,

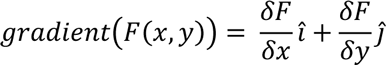

and the inverse tangent was used to return the relative orientation value for each pixel. To assess angular dependence, the pixels with similar orientation assignments were binned to the closest 10 degrees and then the orientation and emission anisotropy values for each pixel were fitted to a cos^2^ function,

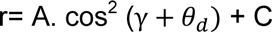

and the absolute amplitude *A* of this wass used as a measure to report the degree of dipole co-alignment.

### Orientation analysis for instantaneous FluoPolScope

Image analysis was performed using custom MATLAB code. Algorithms are available upon request. FAs were segmented using the mApple-tagged paxillin channel as described above. The four polarization resolved quadrants of the integrin-GFP channel were cropped, registered, and throughput-normalized as described in attached related manuscript by Mehta et al. For each FA mask identified from the paxillin channel, background fluorescence was estimated by averaging the dimmest 10 percentile pixels within a 5-pixel wide ring outside of the mask. The 5-pixel wide ring outside of the mask was computed using mathematical morphology. Background corrected polarization resolved intensities (I0, I45, I90, I135) were then summed over each FA. These sum intensities per FA were used to compute ensemble dipole orientation (*o*) and polarization factor (*p*) per FA as follows:

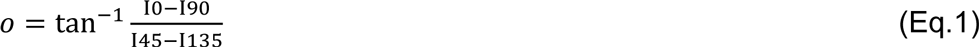

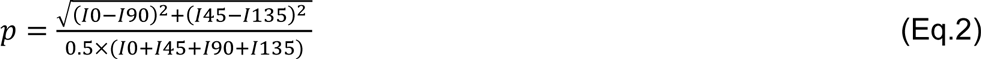

The polarization factor quantifies asymmetry in the angular distribution of dipoles (0=circular distribution, 0.5=elliptical distribution with the major axis twice as long as the minor axis). The ensemble dipole orientation measured by the instantaneous FluoPolScope is independent of the orientation of the FA. Since the ensemble orientation is computed from the ratio of differences of polarization-resolved intensities, it is robust against the isotropic fraction of fluorescence and independent of the total number of fluorophores within a specific FA.

The ensemble orientation of the dipole was measured counter clock-wise from the geometric orientation of the FA. Since the orientation of actin retrograde flow is the same as the orientation of the FA (Fig.3g), the above analysis provides the ensemble orientation of GFP dipoles relative to the actin retrograde flow per FA, as illustrated in Fig. S5b and Fig. 4f for cells expressing αVGFP-constrained and αV-GFP-less-constrained.

### Fitting a Circular Gaussian distribution to experimentally observed GFP dipole orientations

As shown in Fig. 4f, we found that the experimentally measured orientations fit well to the Circular Gaussian distribution (or von Mises distribution), as expected from a contribution of several uncorrelated noise sources.

We defined the Circular Gaussian distribution of orientations by modifying the definition of Circular Gaussian distribution of directions:

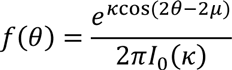

where μ is the mean orientation of the Circular Gaussian, κ is a positive number called the concentration parameter, and *I*_0_(κ) is the modified Bessel function of order 0. The parameter μ and 1/κ are analogous to mean and variance, respectively, of the Gaussian distribution of a linear random variable. The factor 2 in cos (2θ − 2μ) gives rise to a function that has an angular periodicity of π observed in orientation (or axial) data instead of angular periodicity of 2π observed in directional data.

The mean orientation μ and concentration parameter κ were computed using the maximum likelihood estimation functions provided with the CircStat toolbox for MATLAB. The functions were modified to work with orientation data rather than directional data.

### Rosetta simulations

Low energy orientations between the inserted GFP and integrin were efficiently sampled using Rosetta. We treated the integrin and GFP as rigid bodies and carried out Monte Carlo sampling of alternative conformations for the connecting linkers at the integrin-GFP junctions. Backbone torsion angles drawn from short randomly selected polypeptide segments from the protein databank were inserted into the linker regions, and those combinations that enabled the connections at both the N and C-termini of GFP to be closed without steric clashes between GFP and integrin were selected. The geometry at the junctions was then regularized by torsion space energy minimization in the loop regions. Integrin sequences within two residues of the insertion site, linker residues, and GFP residues that vary in position or are disordered in GFP structures (residues 1-5 (MVSKG) and 228-238 (GITLGMDELYK) were considered the loop regions that were subjected to backbone optimization. Only the integrin c-subunit needed to be considered in modeling because the GFP was inserted distal to and never clashed with the β-subunit. The αV model utilized the βV-propeller and thigh domains of PDBID 4GIE^9^.

For each integrin-GFP fusion construct, Rosetta output an ensemble of structures that effectively sampled low energy GFP-integrin orientations. Ensembles were superimposed on the integrin and ranked according to junction loop energy (Fig. S6a). GFP dipole orientation^10,11^ was kindly provided in frame of reference of chain B of PDB ID code 1w7s by the authors of these references as a line with slope x = -0.026, y = 0.871, z = 0.439. A line with this slope drawn through the hydroxyl oxygen atom of the chromophore closely matched a line with α = 6.56 in Fig. 6 of Shi et al.^10^ and was approximated in integrin-GFP ensembles as a line drawn through the N atom of GFP residue Val-112 and the average of the positions of the C atom of Asn-146 and the O atom of Ser-147 in GFP. The integrin frame-of-reference plane was defined by 3 Cα atoms: Asp of the RGD ligand, Arg^4380^ at the junction between the headpiece and the αV leg, and Pro^111^at the junction between the headpiece and the β3 leg. Atoms were chosen for their conserved positions among diverse integrins and integrin conformational states and their proximity to head junctions with flexible domains. This plane was placed into an (x, y, z) frame of reference such that the line defined by the ligand point to the α-leg junction point was nearly co-aligned along the x axis and the line defined by the ligand point to the β leg junction point nearly co-aligned along the the z axis. The specific coordinates were: the ligand was set at the origin (0, 0, 0), the α-leg junction point was set at (54.46, 0, 0), and the β-leg junction point was set at (-0.147, 0, 37.38). The angle of the dipole relative to this integrin plane was denoted as (*ϕ*, θ), in which *ϕ* was rotation around the y axis, and *θ* was rotation around the z axis.

### Prediction of integrin orientation from simulated and experimental GFP orientations

We sought to determine the tilt of the headpiece towards retrograde flow (*ϕ*) assuming that the line connecting α-leg junction point and β-leg junction point was aligned with the retrograde flow/FA long axis. We rotated the Rosetta ensemble of both integrin-GFP chimeras around Y-axis by different angles *ϕ* (Fig. S6 b,c) towards the retrograde flow and computed the overlap between the simulated GFP orientations and experimental GFP orientations as described next.

Rosetta predicts a 3D dipole orientation within the microscope frame of reference, but the instantaneous PolScope measures only the projection of the 3D orientation in the focal plane. Therefore, we projected the 3D dipole orientation of Rosetta models in the focal plane (gray dots, Fig. 4g). Further, Rosetta samples the angular range of the GFP dipole orientations relative to the integrin molecule, but does not predict the probability of occurrence of specific dipole orientations. Therefore, we converted the probability distribution of dipole orientations implied by the circular Gaussian (dashed line in Fig. 4g, Fig. S6, Supplementary Video S1) to the angular range (solid line in the same plots) defined by the full-width-at-half-maximum (FWHM) before computing the overlap of the Rosetta models with experimentally measured dipole orientations. The degree of overlap between projections of simulated dipole orientations and the experimentally measured dipole distribution was computed as follows:

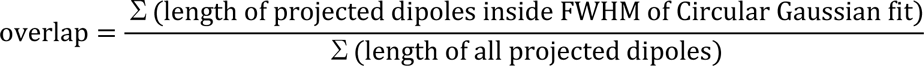

The plot of the overlap (Fig. 4g) for different values of *ϕ* for both αV-GFP-constrained and αV-GFP-less-constrained chimera suggests that the dipole orientations simulated with Rosetta match the best with the experimentally measured orientations when the integrin headpiece is highly tilted towards the retrograde flow relative to the membrane normal.

